# Rapid and direct control of target protein levels with VHL-recruiting dTAG molecules

**DOI:** 10.1101/2020.03.13.980946

**Authors:** Behnam Nabet, Fleur M. Ferguson, Bo Kyung A. Seong, Miljan Kuljanin, Alan L. Leggett, Mikaela L. Mohardt, Amanda Robichaud, Amy S. Conway, Dennis L. Buckley, Joseph D. Mancias, James E. Bradner, Kimberly Stegmaier, Nathanael S. Gray

## Abstract

Chemical biology strategies for directly perturbing protein homeostasis including the degradation tag (dTAG) system provide temporal advantages over genetic approaches and improved selectivity over small molecule inhibitors. We describe dTAG^V^-1, an exclusively selective VHL-recruiting dTAG molecule, to rapidly degrade FKBP12^F36V^-tagged proteins. dTAG^V^-1 overcomes a limitation of previously reported CRBN-recruiting dTAG molecules to degrade recalcitrant oncogenes, supports combination degrader studies and facilitates investigations of protein function in cells and mice.

## INTRODUCTION

Modulating protein abundance with small molecule degraders is a powerful approach for investigating functional consequences of rapid and direct protein loss, without alteration of corresponding mRNA levels. Degraders including heterobifunctional degraders (also known as PROteolysis-TArgeting Chimeras or PROTACs) and non-chimeric molecular glues, co-opt an E3 ubiquitin ligase to induce rapid and reversible proteasome-mediated degradation.^1^ Achieving immediate target protein loss with degraders provides a crucial advantage over genetic knockout or knockdown approaches, which require a significant delay to achieve significant protein reduction.^2^ However, degrader development is hindered by a reliance on target-specific chemical matter, which is unavailable for the majority of the proteome. To address this challenge, several strategies aimed at the direct control of cellular protein levels have been recently developed, including methods that use small molecules^3-9^ or antibodies.^10^

We previously described a versatile approach known as the degradation tag (dTAG) system to rapidly deplete any tagged target protein in cells and in mice.^6^ The dTAG system is a dual component platform requiring the expression of FKBP12^F36V^ in-frame with a gene-of-interest and treatment with a heterobifunctional dTAG molecule (dTAG-13)^11^ that engages FKBP12^F36V^ and cereblon (CRBN), an E3 ubiquitin ligase (Fig. S1a-b). This interaction leads to exclusive degradation of the FKBP12^F36V^-tagged protein. Studies degrading oncoproteins, transcription and chromatin regulators, and kinases exemplify the utility of the dTAG system for drug target validation and discovery.^6, 11-17^ Despite this broad applicability, we observed context- and protein-specific differences in the effectiveness of dTAG-13 for inducing target protein degradation. Here, we report the synthesis, characterization and utility of a second generation, *in vivo*-compatible dTAG molecule that recruits the von Hippel-Lindau (VHL) E3 ligase complex, dTAG^V^-1 (BNN-01-004) (Fig. 1a-b). We demonstrate that dTAG^V^-1 degrades fusion proteins recalcitrant to CRBN-mediated degradation, exemplified by EWS/FLI, a driver of Ewing sarcoma. Collectively, this study describes an important extension to the dTAG platform, towards a universally applicable strategy for direct protein control.

**Fig. 1.**
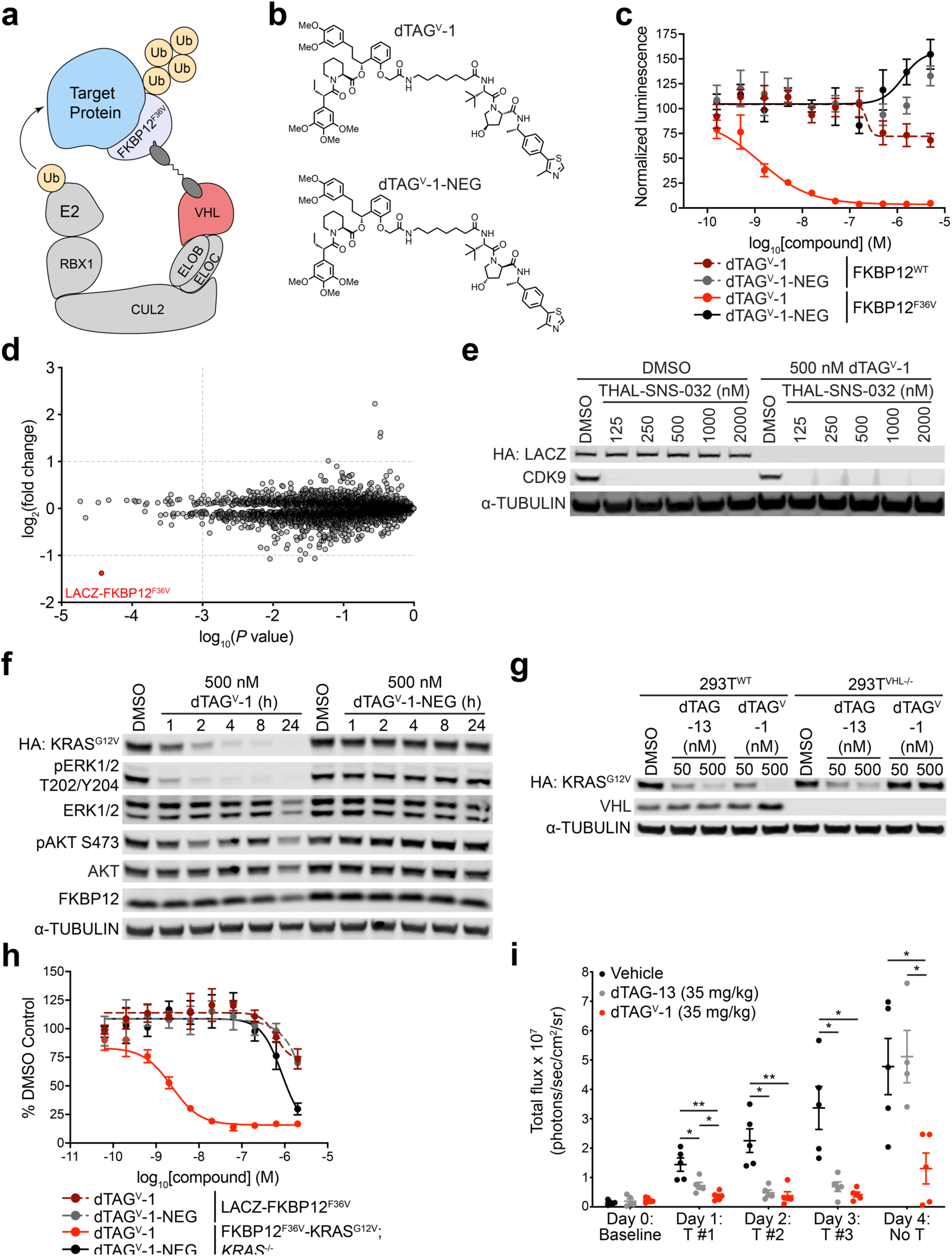
dTAG^V^-1 is an exclusively selective, *in vivo*-compatible degrader of FKBP12^F36V^-tagged proteins. (**a**) Schematic depiction of the dTAG system using VHL-recruiting dTAG molecules. VHL-recruiting dTAG molecules promote ternary complex formation between the FKBP12^F36V^-tagged target protein and E3 ubiquitin ligase complex, inducing target protein ubiquitination and degradation. (**b**) Chemical structures of dTAG^V^-1 and dTAG^V^-1-NEG. (**c**) DMSO-normalized ratio of Nluc/Fluc signal of 293FT FKBP12^WT^-Nluc or FKBP12^F36V^-Nluc cells treated with dTAG^V^-1 or dTAG^V^-1-NEG molecules for 24 h. Data are presented as mean ± s.d. of *n* = 4 biologically independent samples and are representative of *n* = 3 independent experiments. (**d**) Protein abundance after treatment of PATU-8902 LACZ-FKBP12^F36V^ clone with 500 nM dTAG^V^-1 for 4 h compared to DMSO treatment. Volcano plots depict fold change abundance relative to DMSO versus *P* value. Significance designations derived from a permutation-based FDR estimation (q < 0.05) are provided in Supplementary Dataset 1. Data are from *n* = 3 biologically independent samples. (**e**) Immunoblot analysis of PATU-8902 LACZ-FKBP12^F36V^ clone co-treated with DMSO, THAL-SNS-032 and/or dTAG^V^-1 as indicated for 24 h. (**f**) Immunoblot analysis of PATU-8902 FKBP12^F36V^-KRAS^G12V^; *KRAS*^-/-^ clone treated with DMSO, dTAG^V^-1, or dTAG^V^-1-NEG for the indicated time-course. (**g**) Immunoblot analysis of 293T^WT^ or 293T^VHL-/-^ FKBP12^F36V^-KRAS^G12V^ cells treated with DMSO or the indicated dTAG molecules for 24 h. Data in **e**-**g** are representative of *n* = 3 independent experiments. (**h**) DMSO-normalized antiproliferation of PATU-8902 LACZ-FKBP12^F36V^ or FKBP12^F36V^-KRAS^G12V^; *KRAS*^-/-^ clones treated with dTAG^V^-1 or dTAG^V^-1-NEG for 120 h. Cells were cultured as ultra-low adherent 3D-spheroid suspensions. Data are presented as mean ± s.d. of *n* = 4 biologically independent samples and are representative of *n* = 3 independent experiments. (**i**) Bioluminescence imaging to evaluate degradation of luciferase-FKBP12^F36V^ was performed daily as follows: day 0 to establish baseline signal, day 1-3 to monitor luciferase-FKBP12^F36V^ signal 4 h after vehicle or dTAG molecule treatment (T), day 4 to monitor duration of luciferase-FKBP12^F36V^ signal 28 h after third and final vehicle or dTAG molecule treatment. Total flux for each mouse is depicted. Data are presented from vehicle (*n* = 5 biologically independent mice at day 0-4), dTAG-13 (*n* = 5 biologically independent mice at day 0-3; *n* = 4 biologically independent mice at day 4) or dTAG^V^-1 (*n* = 5 biologically independent mice at day 0-4) treated mice. *P* value derived from a two-tailed Welch’s *t*-test (\**P* < 0.05, \*\**P* < 0.01).

## RESULTS

### dTAG^V^-1 is a potent and selective in vivo-compatible degrader

To identify a VHL-recruiting dTAG molecule, we synthesized ortho-AP1867-conjugated analogs with varying VHL-binding ligands and linker composition and screened for cellular activity in 293FT FKBP12^WT^-Nluc and FKBP12^F36V^-Nluc dual luciferase systems^6^ (Fig. 1b-c and Supplementary Fig. 1c-d). dTAG^V^-1 induced potent degradation of FKBP12^F36V^-Nluc with no effects on FKBP12^WT^-Nluc, demonstrating that dTAG^V^-1 is an FKBP12^F36V^-selective degrader (Fig. 1c). dTAG^V^-1-NEG, a diastereomer of dTAG^V^-1 that can no longer bind and recruit VHL^18^ and functions as a heterobifunctional negative control, had no activity on FKBP12^F36V^-Nluc or FKBP12^WT^-Nluc (Fig. 1b-c). These effects were recapitulated in PATU-8902 LACZ-FKBP12^F36V^ cells,^17^ corroborating effective degradation of FKBP12^F36V^-fusions in a pancreatic ductal adenoma carcinoma (PDAC) context (Supplementary Fig. 1e). In both assays, levels of FKBP12^F36V^-fusion degradation with dTAG^V^-1 were comparable to CRBN-recruiting dTAG molecules (dTAG-13 and dTAG-47).^13^ Use of dTAG^V^-1-NEG abrogated these effects comparably to the matched heterobifunctional negative control compounds that cannot bind and recruit CRBN^19, 20^ (dTAG-13-NEG and dTAG-47-NEG) (Supplementary Fig. 1b-e). Multiplexed quantitative mass spectrometry-based proteomics demonstrated that LACZ-FKBP12^F36V^ was the only significantly degraded protein in the proteome (fold change > 2.0, *P* value < 0.001, FDR q < 0.05) upon treatment of PATU-8902 LACZ-FKBP12^F36V^ cells with 500 nM dTAG^V^-1, confirming the exquisite selectivity of the dTAG system (Fig. 1d and Supplementary Dataset 1). No significantly degraded targets were observed with dTAG^V^-1-NEG (Supplementary Fig. 1f and Supplementary Dataset 1).

We next evaluated the utility of dTAG^V^-1 for combinatorial degrader studies. dTAG^V^-1 was effectively combined with THAL-SNS-032, a previously reported CRBN-recruiting CDK9 degrader^21^ (Fig. 1e). Pronounced degradation of LACZ-FKBP12^F36V^ and CDK9 was observed to levels comparable to those with treatment of each degrader alone, avoiding potential substrate competition effects.^22^ To confirm the utility of dTAG^V^-1 for target validation, we evaluated KRAS^G12V^ degradation, an oncogenic driver of PDAC, in PATU-8902 FKBP12^F36V^-KRAS^G12V^; *KRAS*^-/-^ cells.^17^ dTAG^V^-1 treatment led to rapid KRAS^G12V^ degradation, which was rescued by use of dTAG^V^-1-NEG, pre-treatment with proteasome-inhibitor (carfilzomib) or Nedd8 activating enzyme inhibitor (MLN4924), and VHL knockout, consistent with the mechanism of action of a VHL-recruiting degrader (Fig. 1f-g and Supplementary Fig. 2a-b). We also observed the expected collapse in cellular signaling and diminished cell proliferation in 2D-monolayer and 3D-spheroid cultures upon FKBP12^F36V^-KRAS^G12V^ degradation with dTAG^V^-1 treatment, to levels comparable to CRBN-recruiting dTAG molecules (Fig. 1h and Supplementary Fig. 2c-e).

To confirm the *in vivo* applicability of dTAG^V^-1, we characterized the pharmacokinetic (PK) and pharmacodynamic (PD) profile of dTAG^V^-1 in mice. dTAG^V^-1 demonstrated improved properties compared to dTAG-13, with a longer half-life (T_1/2_ = 4.43, 2.41 h respectively) and greater exposure (AUC_inf_ = 18517, 6140 hr*ng mL^-1^ respectively) by intraperitoneal administration at 10 mg/kg (Supplementary Table 1). To report on the PD profile of dTAG molecules, we employed MV4;11 luciferase-FKBP12^F36V^ (luc-FKBP12^F36V^) cells^6^ that allow non-invasive monitoring of bioluminescent signal upon dTAG molecule administration in mice. Following tail vein injection of MV4;11 luc-FKBP12^F36V^ cells and establishment of leukemic burden, we performed daily bioluminescent measurements 4 h after vehicle, 35 mg/kg dTAG-13 or 35 mg/kg dTAG^V^-1 administration. Striking loss of bioluminescent signal was achieved 4 h after the first administration of dTAG^V^-1 (Fig. 1i and Supplementary Fig. 3). Consistent loss of bioluminescent signal was observed 4 h after each of the three dTAG-13 or dTAG^V^-1 administrations. Compared to dTAG-13, improved duration of degradation was also observed with dTAG^V^-1, with degradation evident 28 h after the final administration. These results support the use of dTAG^V^-1 as a potent and selective molecule to evaluate target-specific degradation *in vitro* and *in vivo*.

### dTAG^V^-1 enables evaluation of EWS/FLI degradation in Ewing Sarcoma

There is long-standing interest in the oncology community in identifying targetable dependencies in Ewing sarcoma. Ewing sarcoma is driven by the translocation of *EWSR1* and members of the *ETS* transcription factors, most commonly *FLI*, giving rise to a fusion transcription factor oncoprotein that activates an aberrant transcriptional program.^23^ Currently, there are a shortage of model systems and direct-acting agents that allow modulation of EWS/FLI levels or activity with precise kinetic control. To evaluate the effects of EWS/FLI degradation in Ewing sarcoma, we selected EWS502 cells, which are highly dependent on EWS/FLI for proliferation, and developed FKBP12^F36V^-EWS/FLI; *EWS/FLI*^-/-^ cells (Supplementary Fig. 4a-b). FKBP12^F36V^-GFP was expressed in EWS502 cells as a control and was effectively degraded upon treatment with dTAG^V^-1 and dTAG-13, indicating the ability to co-opt both CRBN and VHL for targeted degradation in this cell line (Fig. 2a). However, FKBP12^F36V^-EWS/FLI was only susceptible to degradation upon treatment with dTAG^V^-1 in EWS502 cells, which was evident within one hour of treatment, highlighting the necessity of comparative evaluation of both CRBN- and VHL-recruiting dTAG molecules (Fig. 2b and Supplementary Fig. 4c). To confirm FKBP12^F36V^-EWS/FLI degradation and identify immediate downstream EWS/FLI targets, we performed acute quantitative mass spectrometry-based proteomics and observed FKBP12^F36V^-EWS/FLI was the most significantly downregulated target protein upon dTAG^V^-1 treatment (Fig. 2c and Supplementary Dataset 1). dTAG^V^-1 led to a decrease in downstream EWS/FLI targets such as NKX2-2 and Gene Set Enrichment Analysis (GSEA) of the downregulated target proteins identified known EWS/FLI and Ewing sarcoma signatures (Fig. 2c-e). Potent antiproliferative effects were also observed upon EWS/FLI degradation at longer time-points, which were absent with dTAG^V^-1-NEG treatment and in FKBP12^F36V^-GFP cell lines (Fig. 2f and Supplementary Fig. 4d). Prior work indicates that BET bromodomain inhibitors and degraders may have applications in Ewing Sarcoma.^24^ To investigate potential synergy between direct and indirect repression of the EWS/FLI transcriptional program, we evaluated degradation of EWS/FLI in combination with BET bromodomain degradation. We observed that dBET6, a CRBN-recruiting, pan-BET bromodomain degrader,^25^ synergized strongly with VHL-mediated EWS/FLI degradation (Supplementary Fig. 5a-c). Together, this data exemplifies the utility of VHL-recruiting dTAG molecules and provides model systems to evaluate the acute and prolonged consequences of EWS/FLI loss.

**Fig. 2.**
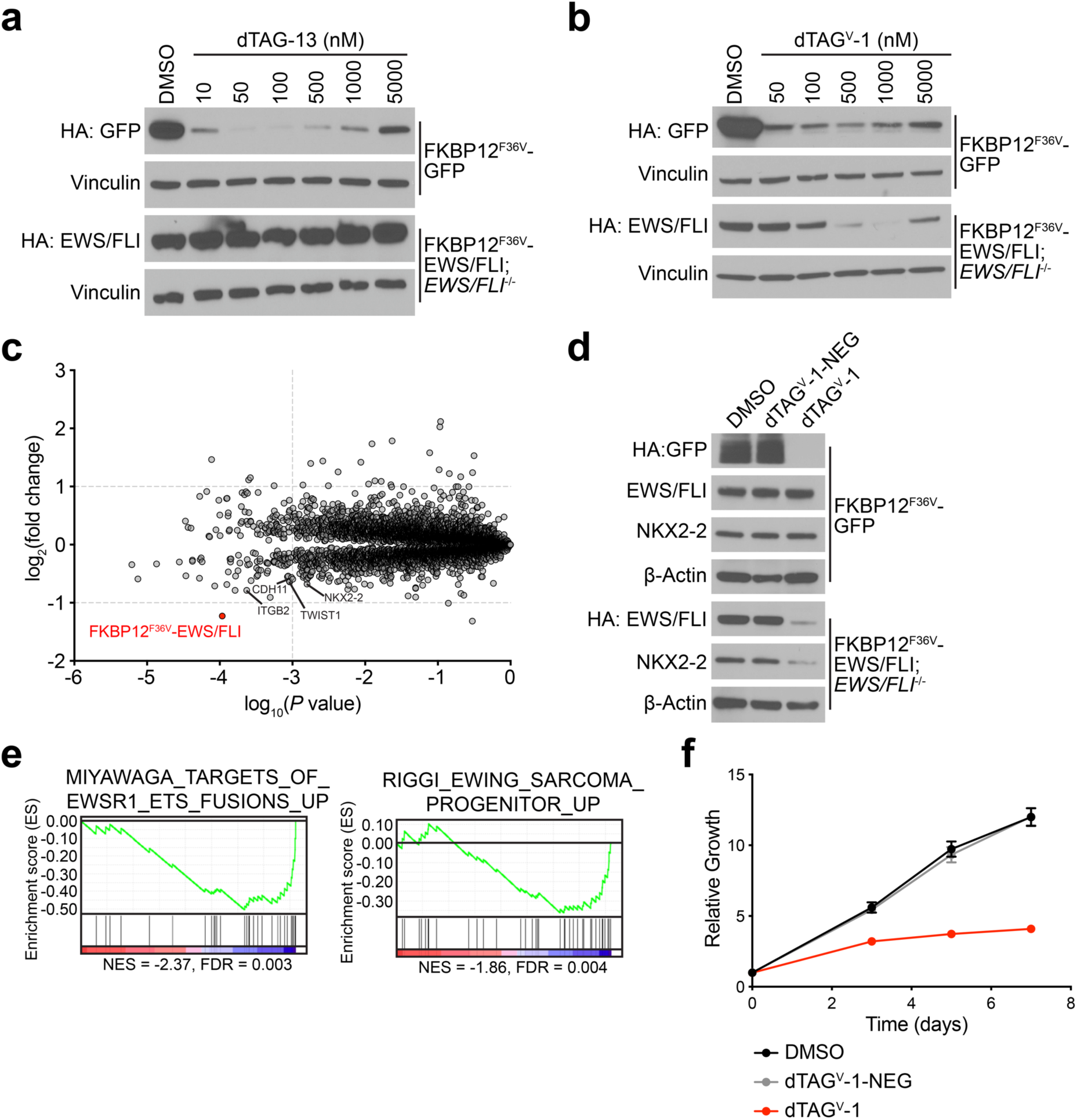
EWS/FLI degradation reverses abnormal proteomic signaling and proliferation. (**a-b**) Immunoblot analysis of EWS502 FKBP12^F36V^-GFP or FKBP12^F36V^-EWS/FLI; *EWS/FLI*^-/-^ cells treated with DMSO (a-b), dTAG-13 (a), or dTAG^V^-1 (b) for 24 h. Data in **a**-**b** are representative of *n* = 3 independent experiments. (**c**) Protein abundance after treatment of EWS502 FKBP12^F36V^-EWS/FLI; *EWS/FLI*^-/-^ cells with dTAG^V^-1 for 6 h compared to DMSO treatment. Volcano plots depict fold change abundance relative to DMSO versus *P* value. Significance designations derived from a permutation-based FDR estimation (q < 0.05) are provided in Supplementary Dataset 1. Data are from *n* = 3 biologically independent samples. (**d**) Immunoblot analysis of EWS502 FKBP12^F36V^-GFP or FKBP12^F36V^-EWS/FLI; *EWS/FLI*^-/-^ cells treated with DMSO, dTAG^V^-1 or dTAG^V^-1-NEG for 24 h. Data are representative of *n* = 3 independent experiments. (**e**) GSEA signatures upon assessment of significantly downregulated target proteins (FDR < 0.05) after treatment of EWS502 FKBP12^F36V^-EWS/FLI; *EWS/FLI*^-/-^ cells as described in **c**. Data are from *n* = 3 biologically independent samples. (**f**) Antiproliferation of EWS502 FKBP12^F36V^-EWS/FLI; *EWS/FLI*^-/-^ cells treated with DMSO, dTAG^V^-1 or dTAG^V^-1-NEG. Y-axis represent luminescence values relative to day 0. Data are presented as mean ± s.d. of *n* = 8 technical replicates and are representative of *n* = 3 independent experiments.

## DISCUSSION

We report dTAG^V^-1, a potent and exclusively selective VHL-recruiting degrader of FKBP12^F36V^-tagged proteins. dTAG^V^-1 displays improved PK/PD properties and serves as an optimized tool for *in vivo* applications. Through evaluation of mutant KRAS degradation in PDAC models, we show that either CRBN or VHL can be co-opted to alleviate the aberrant biology coordinated by this oncoprotein. By contrast, we observed contextual differences in the ability of these E3 ubiquitin ligase complexes to degrade EWS/FLI, supporting use of dTAG^V^-1 for overcoming current limitations of the dTAG system. We demonstrate that VHL-mediated degradation of EWS/FLI rapidly alters downstream target protein expression and leads to pronounced growth defects in Ewing sarcoma cells, providing a powerful model system to investigate immediate consequences of EWS/FLI loss. This data supports that targeting EWS/FLI for degradation with direct-acting heterobifunctional degraders or molecular glues may be a tractable strategy and identifies potential combination strategies with BET bromodomain degraders. Together, the suite of dTAG molecules and paired controls provided in this study will facilitate evaluation of the functional consequences of precise posttranslational protein removal for an expanded target pool. The dTAG system enables rapid modulation of protein abundance and serves as a versatile strategy to determine whether targeted degradation is a promising drug development approach for a given target *in vitro* and *in vivo*.

## METHODS

### Molecule synthesis

#### General methods

Unless otherwise noted, reagents and solvents were obtained from commercial suppliers and were used without further purification. ^1^H NMR spectra were recorded on 500 MHz Bruker Avance III spectrometer, and chemical shifts are reported in parts per million (ppm, δ) downfield from tetramethylsilane (TMS). Coupling constants (J) are reported in Hz. Spin multiplicities are described as s (singlet), br (broad singlet), d (doublet), t (triplet), q (quartet), and m (multiplet). Mass spectra were obtained on a Waters Acquity UPLC. Preparative HPLC was performed on a Waters Sunfire C18 column (19 mm × 50 mm, 5 μM) using a gradient of 15−95% methanol in water containing 0.05% trifluoroacetic acid (TFA) over 22 min (28 min run time) at a flow rate of 20 mL/min. Assayed compounds were isolated and tested as TFA salts. Purities of assayed compounds were in all cases greater than 95%, as determined by reverse-phase HPLC analysis.

#### dTAGV-1 and dTAGV-1-NEG

**Scheme 1.**
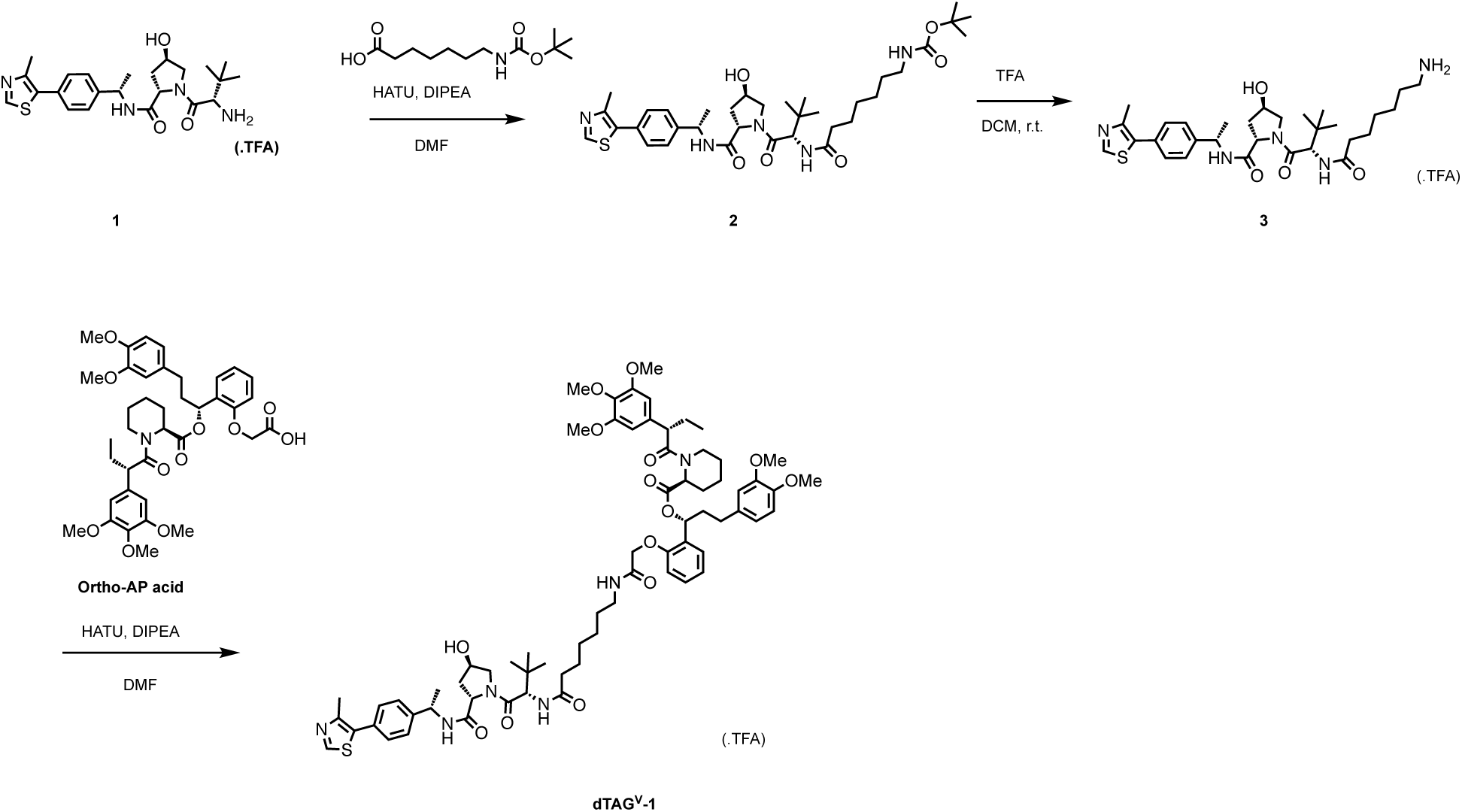
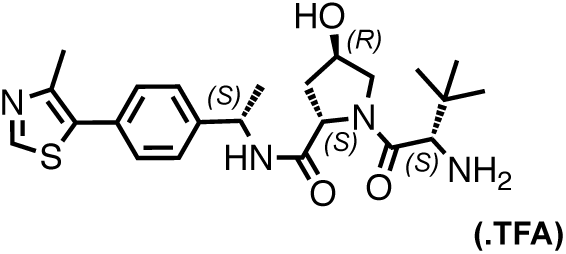

**Chemical structure 1. (2*S***,**4*R*)-1-((*S*)-2-amino-3**,**3-dimethylbutanoyl)-4-hydroxy-*N*-((*S*)-1-(4- (4-methylthiazol-5-yl)phenyl)ethyl)pyrrolidine-2-carboxamide (1)**

The title compound was prepared with minor modifications to the protocol previously described^18^. Specifically trifluoroacetic acid (4N in DCM) was used in place of hydrochloric acid (4N in MeOH) to afford the title compound in quantitative yield.

^1^H NMR (500 MHz, DMSO-*d*_6_) δ 9.00 (s, 1H), 8.60 (dd, *J* = 38.0, 7.8 Hz, 1H), 8.01 (dd, *J* = 16.6, 5.4 Hz, 3H), 7.45 (dd, *J* = 8.4, 2.9 Hz, 2H), 7.39 (dd, *J* = 8.3, 4.9 Hz, 2H), 5.62 (s, 1H), 4.94 (td, *J* = 7.2, 2.6 Hz, 1H), 4.55 (td, *J* = 9.5, 7.5 Hz, 1H), 4.35 (s, 1H), 3.93 (d, *J* = 5.5 Hz, 1H), 3.68 (d, *J* = 11.1 Hz, 1H), 3.51 (dd, *J* = 11.0, 3.8 Hz, 1H), 2.46 (s, 3H), 2.17 – 2.07 (m, 1H), 1.84 – 1.75 (m, 1H), 1.42 – 1.36 (m, 3H), 1.04 (d, *J* = 2.9 Hz, 9H).

LC/MS (ESI^+^): *m/z* 445[M+H^+^]

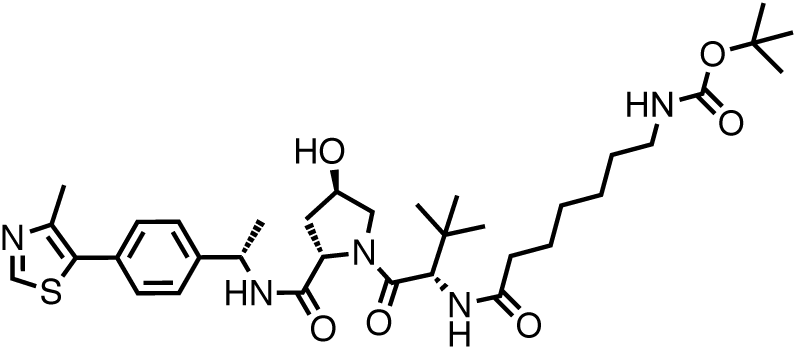

**Chemical structure 2.** *tert***-butyl (7-(((***S***)-1-((2***S*,**4***R***)-4-hydroxy-2-(((***S***)-1-(4-(4-methylthiazol-5-yl)phenyl)ethyl)carbamoyl)pyrrolidin-1-yl)-3**,**3-dimethyl-1-oxobutan-2-yl)amino)-7-oxoheptyl)carbamate (2)**

To a stirred solution of 7-((*tert*-butoxycarbonyl)amino)heptanoic acid (36 mg, 0.12 mmol), HATU (46 mg, 0.12 mmol) and DIPEA (35 μL) in DMF was added **1** (50 mg, 0.1 mmol). The reaction mixture was stirred at room temperature for 16 h. The reaction mixture was diluted with sat. aq. sodium bicarbonate (20 mL) and extracted with EtOAc (3 x 50 mL). The residue was purified by flash chromatography to afford the title compound. (45 mg, 67 %).

^1^H NMR (500 MHz, Methanol-*d*_4_) δ 8.89 (s, 1H), 8.56 (dd, *J* = 8.0, 4.7 Hz, 1H), 8.00 (s, 1H), 7.48 – 7.42 (m, 4H), 6.56 (s, 1H), 5.51 (s, 1H), 5.07 – 4.98 (m, 1H), 4.64 (d, *J* = 8.9 Hz, 1H), 4.59 (dd, *J* = 9.0, 7.7 Hz, 1H), 4.45 (dp, *J* = 4.4, 2.0 Hz, 1H), 3.90 (dt, *J* = 11.2, 1.8 Hz, 1H), 3.80 – 3.74 (m, 2H), 3.05 (d, *J* = 7.0 Hz, 2H), 2.83 (s, 3H), 2.32 – 2.25 (m, 2H), 2.25 – 2.17 (m, 2H), 1.53 (d, *J* = 7.0 Hz, 3H), 1.45 (s, 9H), 1.41 – 1.30 (m, 8H), 1.06 (s, 9H).

LC/MS (ESI^+^): *m/z* 672[M+H^+^]

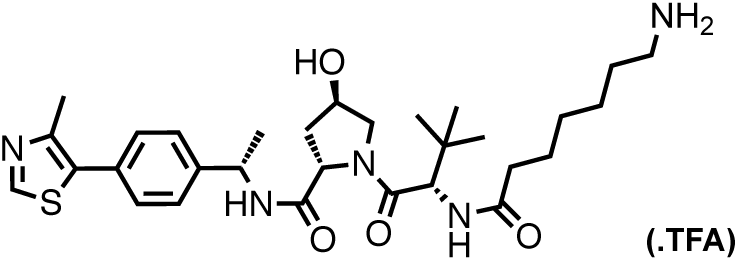

**Chemical structure 3. (2***S*,**4***R***)-1-((***S***)-2-(7-aminoheptanamido)-3**,**3-dimethylbutanoyl)-4-hydroxy-***N***-((***S***)-1-(4-(4-methylthiazol-5-yl)phenyl)ethyl)pyrrolidine-2-carboxamide (3)**

**2** (45 mg, 0.067 mmol) was dissolved in 4N TFA in DCM and stirred at room temperature for 2 h. The reaction mixture was concentrated *in vacuo* to afford the title compound (45 mg, quant), which was used without further purification.

LC/MS (ESI^+^): *m/z* 572[M+H^+^]

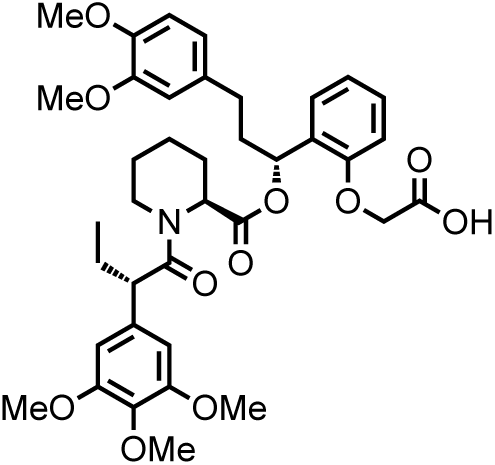

**Chemical structure 4. 2-(2-((***R***)-3-(3**,**4-dimethoxyphenyl)-1-(((***S***)-1-((***S***)-2-(3**,**4**,**5-trimethoxyphenyl)butanoyl)piperidine-2-carbonyl)oxy)propyl)phenoxy)acetic acid (Ortho-AP acid)**

Ortho-AP acid was prepared as previously described.^6^

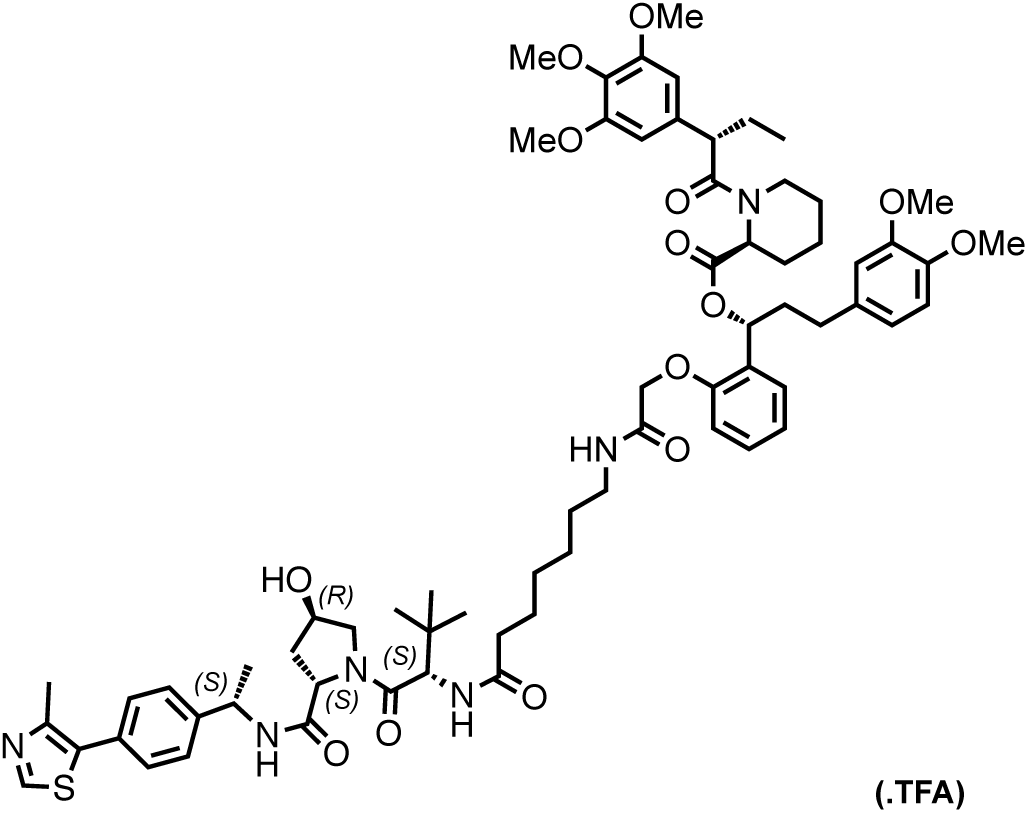

**Chemical structure 5. (***R***)-3-(3**,**4-dimethoxyphenyl)-1-(2-(2-((7-(((***S***)-1-((2***S*,**4***R***)-4-hydroxy-2- (((***S***)-1-(4-(4-methylthiazol-5-yl)phenyl)ethyl)carbamoyl)pyrrolidin-1-yl)-3**,**3-dimethyl-1-oxobutan-2-yl)amino)-7-oxoheptyl)amino)-2-oxoethoxy)phenyl)propyl (***S***)-1-((***S***)-2-(3**,**4**,**5-trimethoxyphenyl)butanoyl)piperidine-2-carboxylate (dTAG**^**V**^**-1)**

To a stirred solution of ortho-AP acid (56 mg, 0.08 mmol), HATU (31 mg, 0.08 mmol), DIPEA (55 μL, 0.2 mmol) in DMF (2 mL) was added **3** (40 mg, 0.067 mmol). The reaction mixture was stirred for 16 h at room temperature, filtered and purified by HPLC to afford the title compound (34 mg, 41 %).

^1^H NMR (500 MHz, DMSO-*d*_6_) δ 8.99 (s, 1H), 8.37 (d, *J* = 7.8 Hz, 1H), 7.77 (dd, *J* = 9.3, 3.3 Hz, 1H), 7.70 (t, *J* = 5.8 Hz, 1H), 7.47 – 7.42 (m, 2H), 7.38 (d, *J* = 8.3 Hz, 2H), 7.21 (ddd, *J* = 8.5, 6.7, 2.4 Hz, 1H), 6.87 (d, *J* = 8.3 Hz, 1H), 6.83 – 6.79 (m, 2H), 6.75 (d, *J* = 2.0 Hz, 1H), 6.64 (dd, *J* = 8.2, 2.0 Hz, 1H), 6.62 (s, 1H), 6.56 (s, 2H), 6.03 (dd, *J* = 8.3, 4.9 Hz, 1H), 5.76 (s, 1H), 5.36 – 5.31 (m, 1H), 5.10 (d, *J* = 3.5 Hz, 1H), 4.92 (p, *J* = 7.3 Hz, 1H), 4.62 – 4.39 (m, 4H), 4.29 (d, *J* = 4.4 Hz, 1H), 4.06 (d, *J* = 13.6 Hz, 1H), 3.91 – 3.84 (m, 1H), 3.75 (s, 1H), 3.72 (s, 3H), 3.71 (s, 2H), 3.70 (s, 3H), 3.64 (d, *J* = 2.6 Hz, 1H), 3.61 (d, *J* = 5.6 Hz, 2H), 3.57 (s, 6H), 3.56 (s, 3H), 3.19 – 3.09 (m, 1H), 3.09 – 3.01 (m, 1H), 2.65 – 2.55 (m, 1H), 2.46 (s, 3H), 2.44 – 2.29 (m, 1H), 2.23 (dt, *J* = 14.5, 7.5 Hz, 1H), 2.16 (d, *J* = 13.2 Hz, 1H), 2.13 – 2.05 (m, 1H), 2.01 (dd, *J* = 12.4, 8.1 Hz, 1H), 1.97 – 1.86 (m, 1H), 1.79 (ddd, *J* = 18.5, 9.4, 5.1 Hz, 1H), 1.60 (qd, *J* = 15.0, 14.5, 9.6 Hz, 2H), 1.49 – 1.40 (m, 1H), 1.38 (d, *J* = 7.0 Hz, 3H), 1.33 (d, *J* = 6.9 Hz, 1H), 1.24 (s, 1H), 1.20 (d, *J* = 17.1 Hz, 6H), 0.94 (s, 9H), 0.81 (t, *J* = 7.3 Hz, 3H).

LC/MS (ESI^+^): *m/z* 1248[M+H^+^], 624[M+H^+^]/2

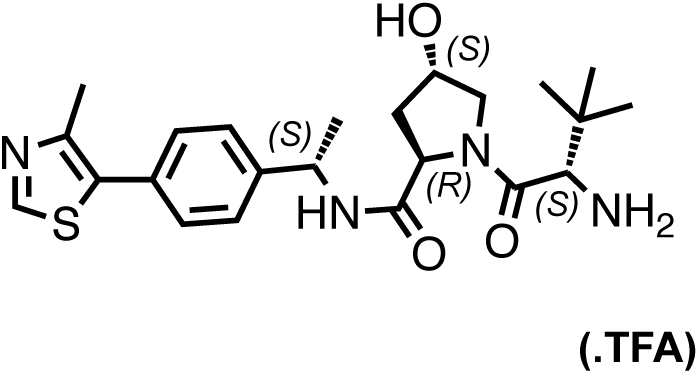

**Chemical structure 6. (2***R*,**4***S***)-1-((***S***)-2-amino-3**,**3-dimethylbutanoyl)-4-hydroxy-***N***-((***S***)-1-(4- (4-methylthiazol-5-yl)phenyl)ethyl)pyrrolidine-2-carboxamide (6)**

The title compound was prepared with minor modifications to the protocol previously described^18^. Specifically trifluoroacetic acid (4N in DCM) was used in place of hydrochloric acid (4N in MeOH) to afford the title compound in quantitative yield.

^1^H NMR (500 MHz, DMSO-*d*_6_) δ 9.00 (s, 1H), 8.42 (d, *J* = 8.0 Hz, 1H), 8.09 (s, 2H), 7.48 – 7.44 (m, 4H), 4.95 (h, *J* = 7.1 Hz, 1H), 4.43 (ddd, *J* = 15.8, 8.5, 5.4 Hz, 2H), 3.92 (q, *J* = 5.5 Hz, 1H), 3.74 (dd, *J* = 10.9, 4.8 Hz, 1H), 3.58 (dd, *J* = 10.8, 3.3 Hz, 1H), 2.47 (s, 3H), 2.10 (ddd, *J* = 12.9, 8.3, 4.4 Hz, 1H), 2.01 – 1.94 (m, 1H), 1.38 (d, *J* = 7.0 Hz, 3H), 1.03 (s, 9H).

LC/MS (ESI^+^): *m/z* 445[M+H^+^]

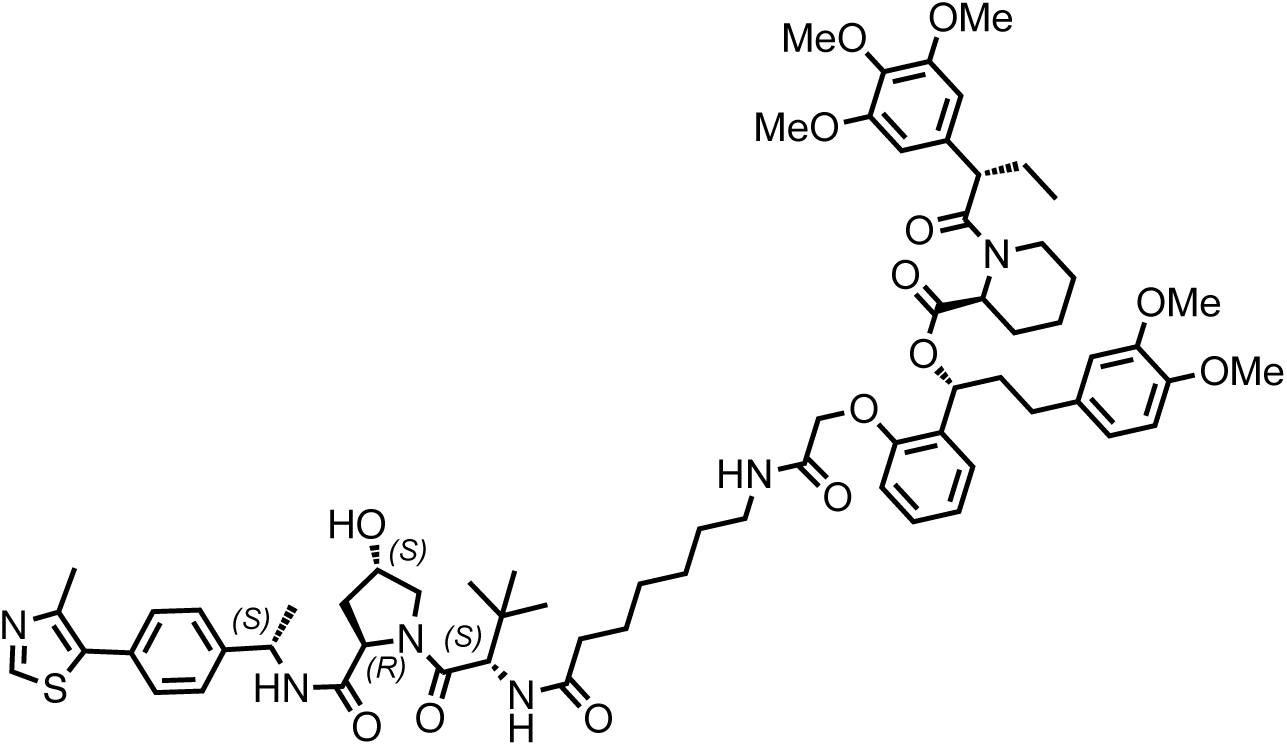

**Chemical structure 7. (*R*)-3-(3**,**4-dimethoxyphenyl)-1-(2-(2-((7-(((*S*)-1-((2*R***,**4*S*)-4-hydroxy-2- (((*S*)-1-(4-(4-methylthiazol-5-yl)phenyl)ethyl)carbamoyl)pyrrolidin-1-yl)-3**,**3-dimethyl-1-oxobutan-2-yl)amino)-7-oxoheptyl)amino)-2-oxoethoxy)phenyl)propyl (*S*)-1-((*S*)-2-(3**,**4**,**5-trimethoxyphenyl)butanoyl)piperidine-2-carboxylate (dTAG**^**V**^**-1-NEG)**

The title compound was prepared from **5** according to scheme 1.

^1^H NMR (500 MHz, DMSO-*d*_6_) δ 8.98 (s, 1H), 8.03 (d, *J* = 8.0 Hz, 1H), 7.89 (d, *J* = 7.8 Hz, 1H), 7.68 (t, *J* = 5.8 Hz, 1H), 7.19 (ddd, *J* = 8.7, 6.6, 2.6 Hz, 1H), 6.86 (d, *J* = 8.3 Hz, 1H), 6.85 – 6.76 (m, 4H), 6.74 (d, *J* = 2.0 Hz, 1H), 6.65 – 6.60 (m, 2H), 6.56 (d, *J* = 2.7 Hz, 2H), 6.03 (dt, *J* = 8.5, 4.6 Hz, 1H), 5.32 (dd, *J* = 5.9, 2.5 Hz, 1H), 4.91 (h, *J* = 7.2, 6.5 Hz, 1H), 4.59 – 4.42 (m, 3H), 4.39 (dd, *J* = 8.0, 5.1 Hz, 2H), 4.31 (p, *J* = 5.2 Hz, 1H), 4.05 (d, *J* = 13.2 Hz, 1H), 3.86 (t, *J* = 7.2 Hz, 1H), 3.81 (dd, *J* = 10.5, 5.4 Hz, 1H), 3.74 (s, 2H), 3.71 (s, 3H), 3.69 (s, 3H), 3.64 (s, 1H), 3.56 (s, 6H), 3.55 (s, 3H), 3.50 (dd, *J* = 10.4, 4.2 Hz, 1H), 3.14 – 3.00 (m, 3H), 2.66 – 2.55 (m, 1H), 2.45 (d, *J* = 1.5 Hz, 3H), 2.43 – 2.30 (m, 1H), 2.24 (dt, *J* = 14.8, 7.7 Hz, 1H), 2.21 – 2.10 (m, 1H), 2.04 (ddd, *J* = 14.5, 8.1, 5.3 Hz, 1H), 2.00 – 1.85 (m, 3H), 1.67 – 1.48 (m, 3H), 1.47 – 1.32 (m, 2H), 1.31 (d, *J* = 7.0 Hz, 3H), 1.27 – 1.00 (m, 7H), 0.97 (s, 9H), 0.81 (t, *J* = 7.3 Hz, 3H).

LC/MS (ESI^+^): *m/z* 1248[M+H^+^], 624[M+H^+^]/2

#### dTAG-13

dTAG-13 was synthesized as previously described.^11^

#### dTAG-47

dTAG-47 was synthesized as previously described.^13^

#### dTAG-13-NEG and dTAG-47-NEG

Compounds were prepared according to the synthetic scheme as previously described.^6, 11, 13^

#### THAL-SNS-032

THAL-SNS-032 was synthesized as previously described.^21^

#### dBET6

dBET6 was synthesized as previously described.^25^

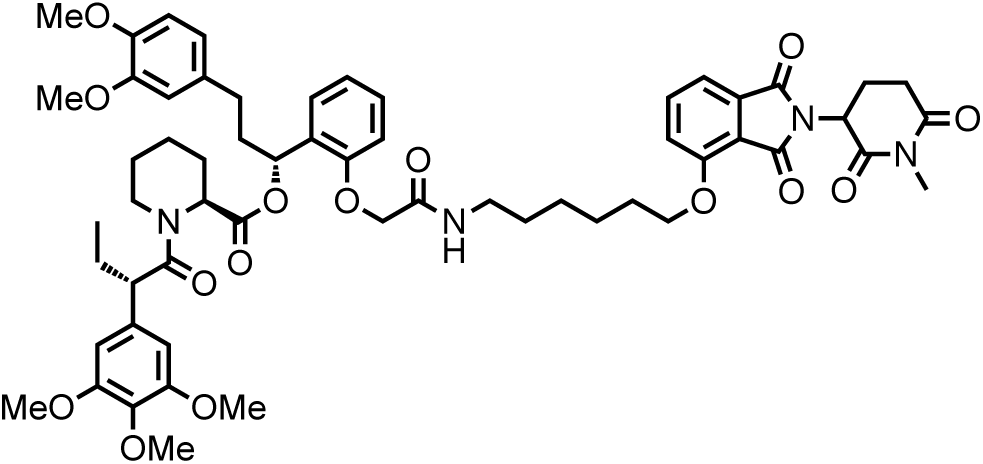

**Chemical structure 8. (1*R*)-3-(3**,**4-dimethoxyphenyl)-1-(2-(2-((6-((2-(1-methyl-2**,**6-dioxopiperidin-3-yl)-1**,**3-dioxoisoindolin-4-yl)oxy)hexyl)amino)-2-oxoethoxy)phenyl)propyl (2*S*)-1-((*S*)-2-(3**,**4**,**5-trimethoxyphenyl)butanoyl)piperidine-2-carboxylate (dTAG-13-NEG)**

^1^H NMR (500 MHz, DMSO-*d*_6_) δ 7.81 (t, *J* = 7.9 Hz, 1H), 7.69 (t, *J* = 5.8 Hz, 1H), 7.50 (dd, *J* = 8.6, 2.2 Hz, 1H), 7.44 (d, *J* = 7.2 Hz, 1H), 7.21 (ddd, *J* = 8.8, 5.8, 3.4 Hz, 1H), 6.88 (d, *J* = 8.3 Hz, 1H), 6.85 – 6.79 (m, 3H), 6.74 (d, *J* = 2.0 Hz, 1H), 6.66 – 6.61 (m, 2H), 6.56 (s, 2H), 6.04 (dd, *J* = 8.3, 4.9 Hz, 1H), 5.36 – 5.30 (m, 1H), 5.15 (dt, *J* = 13.1, 4.4 Hz, 1H), 4.49 (q, *J* = 14.6 Hz, 2H), 4.17 (q, *J* = 6.7 Hz, 2H), 4.05 (d, *J* = 13.3 Hz, 1H), 3.86 (t, *J* = 7.2 Hz, 1H), 3.74 (s, 2H), 3.71 (d, *J* = 1.9 Hz, 5H), 3.69 (s, 3H), 3.64 (s, 1H), 3.57 (s, 6H), 3.55 (s, 3H), 3.08 (q, *J* = 6.7 Hz, 1H), 3.01 (d, *J* = 2.2 Hz, 3H), 2.99 – 2.88 (m, 1H), 2.80 – 2.72 (m, 1H), 2.66 – 2.53 (m, 1H), 2.45 – 2.30 (m, 1H), 2.16 (t, *J* = 8.4 Hz, 2H), 2.08 – 1.85 (m, 3H), 1.70 (h, *J* = 6.2 Hz, 2H), 1.66 – 1.49 (m, 2H), 1.38 (p, *J* = 7.2, 6.7 Hz, 4H), 1.29 – 1.20 (m, 2H), 1.19 – 1.07 (m, 1H), 0.80 (t, *J* = 7.3 Hz, 3H).

MS (ESI) *m/z* 1064 (M + H)^+^. Expected mass from chemical formula C_58_H_70_N_4_O_15_: 1063.21 Da.

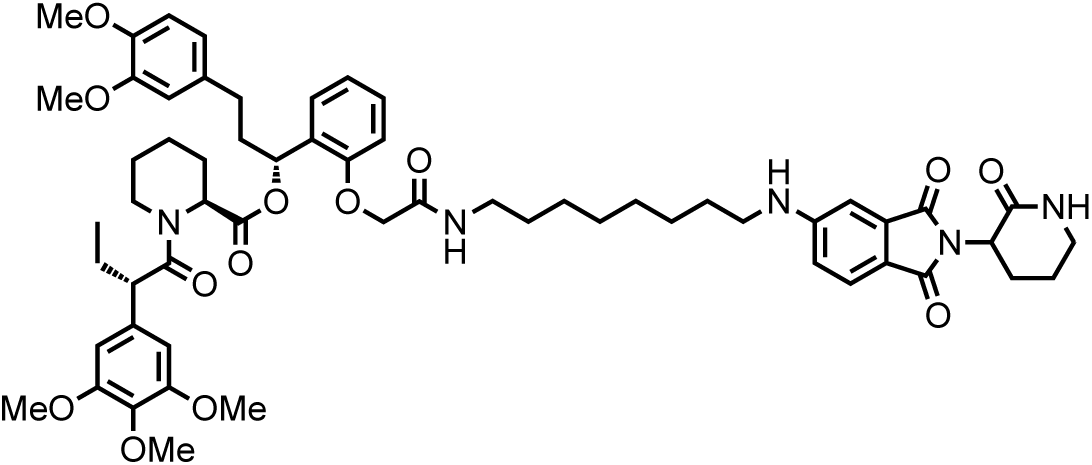

**Chemical structure 9. (1*R*)-3-(3**,**4-dimethoxyphenyl)-1-(2-(2-((8-((1**,**3-dioxo-2-(2-oxopiperidin-3-yl)isoindolin-5-yl)amino)octyl)amino)-2-oxoethoxy)phenyl)propyl (2*S*)-1- ((*S*)-2-(3**,**4**,**5-trimethoxyphenyl)butanoyl)piperidine-2-carboxylate (dTAG-47-NEG)**

^1^H NMR (500 MHz, Methanol-*d*_4_) δ 7.54 (d, *J* = 8.4 Hz, 1H), 7.25 (td, *J* = 7.8, 1.7 Hz, 1H), 6.96 (d, *J* = 2.2 Hz, 1H), 6.91 (t, *J* = 8.1 Hz, 2H), 6.84 (d, *J* = 8.2 Hz, 1H), 6.80 (ddt, *J* = 8.8, 7.4, 1.8 Hz, 2H), 6.76 (d, *J* = 2.0 Hz, 1H), 6.67 (dd, *J* = 8.2, 2.0 Hz, 1H), 6.64 (s, 2H), 6.14 (dd, *J* = 8.2, 6.0 Hz, 1H), 5.42 (d, *J* = 5.4 Hz, 1H), 4.69 (dd, *J* = 11.8, 6.0 Hz, 1H), 4.59 (d, *J* = 15.0 Hz, 1H), 4.44 (d, *J* = 15.0 Hz, 1H), 4.14 (d, *J* = 13.7 Hz, 1H), 3.88 (t, *J* = 7.3 Hz, 1H), 3.82 (d, *J* = 4.7 Hz, 6H), 3.80 (s, 3H), 3.77 (d, *J* = 6.6 Hz, 1H), 3.72 (s, 3H), 3.71 – 3.68 (m, 5H), 3.44 (td, *J* = 11.8, 3.8 Hz, 1H), 3.39 – 3.34 (m, 2H), 3.21 – 3.14 (m, 4H), 2.70 – 2.58 (m, 1H), 2.58 – 2.42 (m, 2H), 2.40 – 2.22 (m, 2H), 2.10 – 2.00 (m, 4H), 1.95 (dt, *J* = 13.9, 6.2 Hz, 1H), 1.74 (dt, *J* = 13.8, 7.0 Hz, 1H), 1.63 (p, *J* = 7.1 Hz, 2H), 1.59 – 1.45 (m, 1H), 1.44 – 1.36 (m, 4H), 1.35 – 0.99 (m, 6H), 0.89 (t, *J* = 7.3 Hz, 3H).

MS (ESI) *m/z* 1063 (M + H)^+^. Expected mass from chemical formula C_59_H_75_N_5_O_13_: 1062.27 Da.

### Cell lines

The following cell lines were employed in this study: 293T (source: Thermo Fisher Scientific, media: DMEM with 10% FBS and 1% Penicillin-Streptomycin), 293FT (source: Thermo Fisher Scientific, media: DMEM with 10% FBS and 1% Penicillin-Streptomycin), PATU-8902 (source: DSMZ, media: DMEM with 10% FBS and 1% Penicillin-Streptomycin), MV4;11 (source: ATCC, media: RPMI with 10% FBS and 1% Penicillin-Streptomycin) and EWS502 (source: Dr. Stephen Lessnick, media: RPMI with 15% FBS and 1% Penicillin-Streptomycin-L-Glutamine). Development of engineered cell lines are detailed below. All cell lines were maintained in 37 °C and 5% CO_2_ incubators and routinely tested negative for mycoplasma contamination using the MycoAlert Kit (Lonza).

### Lentiviral plasmid construction

#### dTAG plasmids

Cloning of pLEX_305-dTAG-KRAS^G12V^ and pLEX_305-LACZ-dTAG were previously described.^6, 17^ To generate pLEX_305-dTAG-GFP and pLEX_305-dTAG-EWS/FLI gateway recombination cloning strategies (Invitrogen) were employed. First, EWS/FLI was cloned into pDONR221 using BP clonase (Invitrogen) after PCR with the following primers containing BP overhangs: Forward-N-E/F-dTAG, 5’- ggggacaagtttgtacaaaaaagcaggcttcgcgtccacggattacagtacct-3’ and Reverse-N-E/F-dTAG, 5’- ggggaccactttgtacaagaaagctgggtcctagtagtagctgcctaagtgtgaaggc-3’. Second, pENTREGFP2 (Addgene #22450) and pDONR221-EWS/FLI were cloned into pLEX_305-N-dTAG using LR clonase (Invitrogen) as previously described.^6^

#### CRISPR/Cas9 plasmids

Cloning of pXPR007-sgGFP and pXPR007-sgKRAS were previously described.^17^ To generate lentiCRISPR v2-Blast-sgFLI_Ex9, sgFLI_Ex9 (5’-GCCTCACGGCGTGCAGGAAG-3’) was cloned into lentiCRISPR v2-Blast vector (Addgene, #83480) using BsmbI restriction sites. The PAM motif of sgFLI_Ex9 is present in an intron, enabling cutting of the endogenous locus only with cDNA rescue.

### Lentivirus production, transduction and cell line development

#### PATU-8902 LACZ-FKBP12^F36V^ and FKBP12^F36V^-KRAS^G12V^; KRAS^-/-^ cells

Development and characterization of PATU-8902 LACZ-FKBP12^F36V^ and FKBP12^F36V^-KRAS^G12V^; *KRAS*^-/-^ clones were previously described.^17^

#### 293T^WT^ FKBP12^F36V^-KRAS^G12V^ and 293T^VHL-/-^ FKBP12^F36V^-KRAS^G12V^ cells

pLEX305-dTAG-KRAS^G12V^ lentivirus was prepared and concentrated as previously described.^6^ To generate 293T^WT^ FKBP12^F36V^-KRAS^G12V^ and 293T^VHL-/-^ FKBP12^F36V^-KRAS^G12V^ cells, concentrated lentiviral supernatants were applied in the presence of 4 µg/mL polybrene and transduced cell lines were selected with 2 µg/mL puromycin.

#### EWS502 FKBP12^F36V^-GFP and FKBP12^F36V^-EWS/FLI; EWS/FLI ^-/-^ cells

To generate EWS502 FKBP12^F36V^-GFP cells, EWS502 cells were transduced with pLEX_305-dTAG-GFP lentiviral supernatant and selected with puromycin. To generate EWS502 FKBP12^F36V^-EWS/FLI; *EWS/FLI*^-/-^ cells, EWS502 cells were co-transduced with pLEX_305-dTAG-EWS/FLI and lentiCRISPR v2-Blast-sgFLI_Ex9 lentiviral supernatants. Cells were then selected with both puromycin (pLEX_305-dTAG-EWS/FLI) and blasticidin (lentiCRISPR v2-Blast-sgFLI_Ex9). Knockout of endogenous EWS/FLI and the expression of exogenous FKBP12^F36V^-EWS/FLI were confirmed by immunoblot and pooled transduced cell populations were employed for this study.

### FKBP12^WT^ and FKBP12^F36V^ dual luciferase assay

Dual luciferase assays were performed using 293FT FKBP12^WT^-Nluc and FKBP12^F36V^-Nluc cells as previously described^6^ with the following modifications. Cells were plated at 2000 cells per well in 20 µL of appropriate media in 384-well white culture plates (Corning), allowed to adhere overnight, and 100 nL of compounds were added using a Janus Workstation pin tool (PerkinElmer) for 24 hours at 37 °C. To evaluate Fluc signal, plates were brought to room temperature, 20 µL of Dual-Glo Reagent (Promega) was added for 10 minutes and luminescence was measured on an Envision 2104 plate reader (PerkinElmer). Subsequently, 20 µL of Dual-Glo Stop & Glo Reagent (Promega) was added for 10 minutes and luminescence was again measured to capture Nluc signal. Data were analyzed as previously described.^6^

### Immunoblotting

PATU-8902 LACZ-FKBP12^F36V^ and FKBP12^F36V^-KRAS^G12V^; *KRAS*^-/-^ cells and 293T cells were lysed as previously described^6, 17^ and immunoblotting was performed using an Odyssey CLx Imager (LI-COR) as previously described.^6, 17^ EWS502 cells were lysed with Cell Lysis Buffer (Cell Signaling Technology) supplemented with cOmplete, EDTA-free Protease Inhibitor Cocktail (Roche). Proteins were separated by SDS-PAGE and transferred to PVDF membranes, which were blocked with 5% milk in TBS-T and incubated with primary antibodies overnight. Membranes were washed 5 times with TBS-T and incubated with the appropriate horseradish peroxidase-conjugated secondary antibodies. The following primary antibodies were employed in this study: HA (Cell Signaling, #3724 and #2367), phospho-ERK1/2 T202/Y204 (Cell Signaling, #4370), ERK1/2 (Cell Signaling, #4696), phospho-AKT S473 (Cell Signaling, #4060), AKT (Cell Signaling, #2920), FKBP12 (Abcam, #ab24373), GFP (Cell Signaling, #2555), FLI (Abcam, #15289), NKX2-2 (Abcam, #187375), GAPDH (Cell Signaling, #2118), β-Actin (Cell Signaling, #58169), and α-Tubulin (Cell Signaling, #3873). Species-specific fluorescently labelled infrared (IRDye) and peroxidase-linked (Thermo Fisher Scientific) secondary antibodies were employed as appropriate.

### Analysis of cell viability

Cell viability was assayed in 2D-adherent and ultra-low adherent 3D-spheroids using CellTiter-Glo (Promega) as previously described.^6, 17^ Synergy assessments were performed using CellTiter-Glo (Promega) as previously described^26^ with the following modifications. EWS502 cells were plated at 1000 cells per well in 50 µL of appropriate media in 384-well white culture plates (Corning). Luminescence was measured on an Envision 2104 plate reader (PerkinElmer) and Fluostar Omega Reader (BMG Labtech).

### Quantitative proteomics

#### Materials

The following reagents were employed: Isobaric TMT reagents (Thermo Fisher Scientific), BCA protein concentration assay kit (Thermo Fisher Scientific), Empore-C18 material for in-house made StageTips (3 M), Sep-Pak cartridges (100 mg, Waters), solvents for Liquid chromatography (LC) (J.T. Baker), mass spectrometry (MS)-grade trypsin (Thermo Fisher Scientific), Lys-C protease (Wako), and cOmplete protease inhibitors (Millipore Sigma). Unless otherwise noted, all other chemicals were purchased from Thermo Fisher Scientific.

#### MS sample processing

Cell pellets from PATU-8902 and EWS502 cells were lysed using 8 M urea, 200 mM 4-(2-hydroxyethyl)-1-piperazinepropanesulfonic acid (EPPS) at pH 8.5 with protease inhibitors. Samples were further homogenized and DNA was sheered via sonication using a probe sonicator (20× 0.5 sec pulses; level 3). Total protein was determined using a BCA assay and stored at −80°C until future use. A total of 100 µg of protein was aliquoted for each condition and TMT channel for further downstream processing. Protein extracts were reduced using 10 mM dithiothreitol (DTT) for 30 min at room temperature. Next samples were alkylated with 20 mM iodoacetamide for 45 min in the dark at room temperature. To facilitate the removal of incompatible reagents, proteins were precipitated using chloroform methanol. Briefly, to 100 µL of each sample, 400 µL of methanol was added, followed by 100 µL of chloroform with thorough vortexing. Next, 300 µL of HPLC grade water was added and samples were vortexed thoroughly. Each sample was centrifuged at 14,000 xg for 5 min at room temperature. The upper aqueous layer was removed and the protein pellet was washed twice with methanol and centrifuged at 14,000 xg for 5 min at room temperature. Protein pellets were re-solubilized in 200 mM EPPS buffer and digested overnight with Lys-C (1:100, enzyme: protein ratio) at room temperature. The next day, trypsin (1:100 ratio) was added and incubated at 37 °C for an additional 6 h in a ThermoMixer set to 1000 RPM.

To each digested sample, 30% anhydrous acetonitrile was added and 100 µg of peptides were labeled using 200 µg of TMT reagent (TMT1-TMT11). To equalized protein loading, a ratio check was performed by pooling 2 µg of each TMT-labeled sample. Samples were pooled and desalted using in-house packed C18 StageTips and analyzed by LC-MS/MS. Normalization factors were calculated from this label check, samples were mixed 1:1 across all TMT channels and desalted using a 100 mg Sep-Pak solid phase extraction cartridge. Eluted pooled peptides were further fractionated with basic-pH reverse-phase (bRP) HPLC using an Agilent 300 extend C18 column and collected into a 96 deep-well plate. Samples were consolidated into 24 fractions as previously described, and 12 nonadjacent fraction were desalted using StageTips prior to analyses using LC-MS/MS.^27-29^

#### MS data acquisition

All mass spectrometry data was acquired using an Orbitrap Fusion mass spectrometer in-line with a Proxeon NanoLC-1000 UHPLC system. Peptides were separated using an in-house 100 µm capillary column packed with 40 cm of Accucore 150 resin (2.6 um, 150 Å) (Thermo Fisher Scientific) using a 180 min LC gradient per fraction. Eluted peptides were acquired using synchronous precursor selection (SPS-MS3) method for TMT quantification as previously described.^30^ Briefly, MS1 spectra were acquired at 120K resolving power with a maximum of 50 ms in the Orbitrap. MS2 spectra were acquired by selecting the top 10 most abundant features via collisional induced dissociation (CID) in the ion trap using an automatic gain control (AGC) of 15K, quadrupole isolation width of 0.7 m/z and a maximum ion time of 100 ms. For MS3 acquisition, a synchronous precursor selection of 10 fragment ions was acquired with an AGC of 150K for 150 ms and a normalized collision energy of 55.

#### MS data analysis

All acquired data were processed using SEQUEST^31^ and a previously described in-house informatics pipeline.^32-34^ Briefly, peptide spectral libraries were first filtered to a peptide false discovery rate (FDR) of less than 1 % using linear discriminant analysis employing a target decoy strategy. Spectral searches were done using a custom fasta formatted database which included custom sequences for LACZ-FKBP12^F36V^ and FKBP12^F36V^-EWS/FLI, common contaminants, reversed sequences (Uniprot Human, 2014) and the following parameters: 50 ppm precursor tolerance, fully tryptic peptides, fragment ion tolerance of 0.9 Da and a static modification of TMT (+229.163 Da) on lysine and peptide N-termini, carbamidomethylation of cysteine residues (+57.021 Da) were set as static modifications, while oxidation of methionine residues (+15.995 Da) was set as a variable modification. Resulting peptides were further filtered to obtain a 1 % protein FDR and proteins were collapsed into groups. Reporter ion intensities were adjusted to correct for impurities during synthesis of different TMT reagents according to the manufacturer’s specifications. For quantitation, a total sum signal-to-noise of all report ions of 200 was required for analysis. Protein quantitative values were normalized so that the sum of the signal for all protein in each channel was equal to account for sample loading. Gene Set Enrichment Analysis (GSEA)^35^ was performed as indicated in the figure legends.

### Animal studies

#### Compound formulation

For IP injections, dTAG-13 and dTAG^V^-1 were formulated by dissolving into DMSO and then diluting with 20% solutol (Sigma): 0.9 % sterile saline (Moltox) (w:v) with the final formulation containing 5 % DMSO. Maximal solubility of 35 mg/kg and 40 mg/kg were observed for dTAG-13 and dTAG^V^-1, respectively. Formulations were stable at room temperature for 7 days. For IV injections, dTAG-13 and dTAG^V^-1 were formulated by dissolving into DMSO and then diluting with 5 % solutol (Sigma): 0.9 % sterile saline (Moltox) (w:v) with the final formulation containing 5 % DMSO.

#### Pharmacokinetic (PK) evaluation

PK was assessed in 8-week-old C57BL/6J male mice (Jackson Laboratory, #000664) with blood collected at 0.08, 0.25, 0.5, 1, 2, 4, 6, and 8 h (2 mg/kg dTAG-13 intravenous (IV) tail vein, 10 mg/kg dTAG-13 intraperitoneal (IP), and 2 mg/kg dTAG^V^-1 IV tail vein administrations) and 0.25, 0.5, 1, 2, 4, 6, 8, 24 and 48 h (2 mg/kg dTAG^V^-1 IP and 10 mg/kg dTAG^V^-1 IP administrations). Plasma was generated by centrifugation and plasma concentrations were determined by LC-MS/MS following the mass transition 49600à340 AMU. PK parameters were calculated using Phoenix WinNonlin to determine peak plasma concentration (C_max_), oral bioavailability (% F), exposure (AUC), half-life (t_1/2_), clearance (CL), and volume of distribution (V_d_). All procedures were approved by and performed in accordance with standards of the Institute Animal Care and Use Committee (IACUC) at Scripps Florida.

#### Pharmacodynamic (PD) evaluation

Evaluation of degradation of luciferase was performed as previously described.^6^ In brief, 2.5 × 10^5^ viable MV4;11 luc-FKBP12^F36V^ cells were transplanted by tail-vein injection in 8-week-old immunocompromised female mice (NOD.Cg-*Prkdc*^*scid*^*Il2rg*^*tm1Wjl*^/SzJ, NSG; Jackson Laboratory, #005557). Bioluminescence measurements were used to monitor engraftment and establish baseline signal. For treatments, compounds were formulated as described above, administered via IP injection and bioluminescence measurements were performed daily as described in Fig. 1i. All procedures were approved by and performed in accordance with standards of the IACUC at Dana-Farber Cancer Institute.

### Statistical analysis

Information regarding center values, error bars, number of replicates or samples, number of independent experiments, and statistical analyses are described in the corresponding figure and table legends. Experiments were not blinded nor randomized, and sample sizes were not predetermined using statistical analyses.

## DATA AVAILABILITY STATEMENT

Mass spectrometry-based proteomics data files are provided in Supplementary Dataset 1.

## REAGENT AVAILABILITY

Reagents are freely available at: http://graylab.dana-farber.org/probes.html.

## ACKNOWLEDGEMENTS

We thank M. Kostic, N. Kwiatkowski and G. Winter for critical reading of the manuscript and S. Nabet and members of the Gray laboratory for helpful discussions. We gratefully acknowledge S. Gygi for use of CORE for mass spectrometry data analysis software. This work was supported by: American Cancer Society Postdoctoral Fellowship PF-17-010-01-CDD (B.N.), Claudia Adams Barr Program in Innovative Basic Cancer Research Award (B.N.), Katherine L. and Steven C. Pinard Research Fund (B.N. and N.S.G.), Damon Runyon Cancer Research Foundation DRG-2196-14 (D.L.B.), Burroughs Wellcome Fund Career Award in Medical Sciences Award (J.D.M.), NCI U54 CA231637 (K.S.), NCI R01 CA204915 (K.S.), and Hale Center for Pancreatic Cancer Research (J.D.M. and N.S.G.).

## AUTHOR CONTRIBUTIONS

B.N. and F.M.F. conceived and led the study. B.N., F.M.F, and D.L.B. designed and performed molecule synthesis. B.N. and A.L.L. performed the FKBP12 dual luciferase experiments. B.N., A.L.L., and M.L.M. designed and performed the LACZ and KRAS studies. B.K.S. designed and developed the EWS/FLI systems and B.N., B.K.S. and M.L.M. performed experiments using these systems. B.N. and B.K.S. prepared samples for proteomics studies and M.K. performed the mass spectrometry assessment and analyses. B.N., B.K.S., A.R., and A.S.C. designed and performed mouse studies. J.D.M., J.E.B., K.S. and N.S.G. provided technical advice, data interpretation and supervised the study. B.N. and F.M.F. wrote the manuscript with input from all authors.

## COMPETING FINANCIAL INTERESTS

The authors claim the following competing financial interests: B.N., D.L.B., and J.E.B. are inventors on patent applications related to the dTAG system (WO/2017/024318, WO/2017/024319, WO/2018/148440 and WO/2018/148443). The molecules disclosed in this manuscript are the subject of a patent application filed by Dana-Farber Cancer Institute. D.L.B. is now an employee of Novartis. J.E.B. is a Scientific Founder of Syros Pharmaceuticals, SHAPE Pharmaceuticals, Acetylon Pharmaceuticals, Tensha Therapeutics (now Roche), and C4 Therapeutics and is the inventor on IP licensed to these entities. J.E.B. is now an executive and shareholder in Novartis AG. K.S. has previously consulted for Novartis and Rigel Pharmaceuticals and has received research funding from Novartis. N.S.G. is a Scientific Founder and member of the Scientific Advisory Board (SAB) of C4 Therapeutics, Syros, Soltego, Gatekeeper and Petra Pharmaceuticals and has received research funding from Novartis, Astellas, Taiho and Deerfield.

## REFERENCES

1. Lai, A.C. & Crews, C.M. Induced protein degradation: an emerging drug discovery paradigm. Nat Rev Drug Discov 16, 101–114 (2017).

2. Yesbolatova, A., Tominari, Y. & Kanemaki, M.T. Ligand-induced genetic degradation as a tool for target validation. Drug Discov Today Technol 31, 91–98 (2019).

3. Bonger, K.M., Chen, L.C., Liu, C.W. & Wandless, T.J. Small-molecule displacement of a cryptic degron causes conditional protein degradation. Nat. Chem. Biol. 7, 531–537 (2011).

4. Buckley, D.L. et al. HaloPROTACS: Use of Small Molecule PROTACs to Induce Degradation of HaloTag Fusion Proteins. ACS Chem. Biol. 10, 1831–1837 (2015).

5. Chung, H.K. et al. Tunable and reversible drug control of protein production via a self-excising degron. Nat. Chem. Biol. 11, 713–720 (2015).

6. Nabet, B. et al. The dTAG system for immediate and target-specific protein degradation. Nat Chem Biol 14, 431–441 (2018).

7. Nishimura, K., Fukagawa, T., Takisawa, H., Kakimoto, T. & Kanemaki, M. An auxin-based degron system for the rapid depletion of proteins in nonplant cells. Nature methods 6, 917–922 (2009).

8. Koduri, V. et al. Peptidic degron for IMiD-induced degradation of heterologous proteins. Proc. Natl. Acad. Sci. U. S. A. 116, 2539–2544 (2019).

9. Neklesa, T.K. et al. Small-molecule hydrophobic tagging-induced degradation of HaloTag fusion proteins. Nat. Chem. Biol. 7, 538–543 (2011).

10. Clift, D., So, C., McEwan, W.A., James, L.C. & Schuh, M. Acute and rapid degradation of endogenous proteins by Trim-Away. Nat. Protoc. 13, 2149–2175 (2018).

11. Erb, M.A. et al. Transcription control by the ENL YEATS domain in acute leukaemia. Nature 543, 270–274 (2017).

12. Brunetti, L. et al. Mutant NPM1 Maintains the Leukemic State through HOX Expression. Cancer Cell 34, 499–512 e499 (2018).

13. Huang, H.T. et al. MELK is not necessary for the proliferation of basal-like breast cancer cells. Elife 6, e26693 (2017).

14. Weintraub, A.S. et al. YY1 Is a Structural Regulator of Enhancer-Promoter Loops. Cell 171, 1573–1588 e1528 (2017).

15. Boija, A. et al. Transcription Factors Activate Genes through the Phase-Separation Capacity of Their Activation Domains. Cell 175, 1842–1855 e1816 (2018).

16. Weissmiller, A.M. et al. Inhibition of MYC by the SMARCB1 tumor suppressor. Nat Commun 10, 2014 (2019).

17. Ferguson, F.M. et al. Discovery of a selective inhibitor of Doublecortin Like Kinase 1. Nat. Chem. Biol. In Press (2020).

18. Raina, K. et al. PROTAC-induced BET protein degradation as a therapy for castration-resistant prostate cancer. Proc. Natl. Acad. Sci. U. S. A. 113, 7124–7129 (2016).

19. Brand, M. et al. Homolog-Selective Degradation as a Strategy to Probe the Function of CDK6 in AML. Cell Chem Biol 26, 300–306 e309 (2019).

20. Huang, H.T. et al. A Chemoproteomic Approach to Query the Degradable Kinome Using a Multi-kinase Degrader. Cell Chem Biol 25, 88–99 e86 (2018).

21. Olson, C.M. et al. Pharmacological perturbation of CDK9 using selective CDK9 inhibition or degradation. Nat. Chem. Biol. 14, 163–170 (2018).

22. Sperling, A.S. et al. Patterns of substrate affinity, competition, and degradation kinetics underlie biological activity of thalidomide analogs. Blood 134, 160–170 (2019).

23. Riggi, N. et al. EWS-FLI1 utilizes divergent chromatin remodeling mechanisms to directly activate or repress enhancer elements in Ewing sarcoma. Cancer Cell 26, 668–681 (2014).

24. Gollavilli, P.N. et al. EWS/ETS-Driven Ewing Sarcoma Requires BET Bromodomain Proteins. Cancer Res. 78, 4760–4773 (2018).

25. Winter, G.E. et al. BET Bromodomain Proteins Function as Master Transcription Elongation Factors Independent of CDK9 Recruitment. Mol. Cell 67, 5–18 e19 (2017).

26. Li, Z. et al. Development and Characterization of a Wee1 Kinase Degrader. Cell Chem Biol (2019).

27. Navarrete-Perea, J., Yu, Q., Gygi, S.P. & Paulo, J.A. Streamlined Tandem Mass Tag (SL-TMT) Protocol: An Efficient Strategy for Quantitative (Phospho)proteome Profiling Using Tandem Mass Tag-Synchronous Precursor Selection-MS3. J Proteome Res 17, 2226–2236 (2018).

28. Paulo, J.A. Nicotine alters the proteome of two human pancreatic duct cell lines. JOP 15, 465–474 (2014).

29. Paulo, J.A. & Gygi, S.P. Nicotine-induced protein expression profiling reveals mutually altered proteins across four human cell lines. Proteomics 17 (2017).

30. McAlister, G.C. et al. MultiNotch MS3 enables accurate, sensitive, and multiplexed detection of differential expression across cancer cell line proteomes. Anal Chem 86, 7150–7158 (2014).

31. Eng, J.K., McCormack, A.L. & Yates, J.R. An approach to correlate tandem mass spectral data of peptides with amino acid sequences in a protein database. J Am Soc Mass Spectrom 5, 976–989 (1994).

32. Beausoleil, S.A., Villen, J., Gerber, S.A., Rush, J. & Gygi, S.P. A probability-based approach for high-throughput protein phosphorylation analysis and site localization. Nat Biotechnol 24, 1285–1292 (2006).

33. Elias, J.E. & Gygi, S.P. Target-decoy search strategy for increased confidence in large-scale protein identifications by mass spectrometry. Nat Methods 4, 207–214 (2007).

34. Huttlin, E.L. et al. A tissue-specific atlas of mouse protein phosphorylation and expression. Cell 143, 1174–1189 (2010).

35. Subramanian, A. et al. Gene set enrichment analysis: a knowledge-based approach for interpreting genome-wide expression profiles. Proc. Natl. Acad. Sci. U. S. A. 102, 15545–15550 (2005).

