## Supplementary Tables and Figures for "Rapid and direct control of target protein levels with VHL-recruiting dTAG molecules"

[1]Department of Cancer Biology, Dana-Farber Cancer Institute, Boston, Massachusetts, USA.
[2]Department of Biological Chemistry and Molecular Pharmacology, Harvard Medical School, Boston, Massachusetts, USA.
[3]Department of Pediatric Oncology, Dana-Farber Cancer Institute, Boston, Massachusetts, USA.
[4]The Broad Institute of MIT and Harvard, Cambridge, Massachusetts, USA.
[5]Division of Radiation and Genome Stability, Department of Radiation Oncology, Dana-Farber Cancer Institute, Boston, Massachusetts, USA.
[6]Department of Medical Oncology, Dana-Farber Cancer Institute, Boston, Massachusetts, USA.
[7]Department of Medicine, Harvard Medical School, Boston, Massachusetts, USA.
[8]Division of Pediatric Hematology/Oncology, Boston Children's Hospital, Boston, Massachusetts, USA.
[9]Present address: Novartis Institutes for BioMedical Research, Cambridge, Massachusetts, USA.
[10]These authors contributed equally to this work.
*Corresponding author

Correspondence may be addressed to:

Behnam Nabet


Nathanael S. Gray


### SUPPLEMENTARY TABLES

| | | dTAG-13 | | $dTAG^V$-1 | | |
|---|---|---|---|---|---|---|
| **Parameter** | **Unit** | **IV** | **IP** | **IV** | **IP** | **IP** |
| Dose | mg $kg^{-1}$ | 2 | 10 | 2 | 2 | 10 |
| $T_{max}$ | hr | 0.08 | 2.00 | 0.08 | 1.67 | 2.00 |
| $T_{1/2}$ | hr | 1.46 | 2.41 | 3.02 | 3.64 | 4.43 |
| $C_{max}$ | ng $mL^{-1}$ | 2373 | 1263 | 7780 | 595 | 2123 |
| $AUC_{last}$ | hr*ng $mL^{-1}$ | 1242 | 5619 | 3245 | 2245 | 18088 |
| $AUC_{inf}$ | hr*ng $mL^{-1}$ | 1253 | 6140 | 3329 | 3136 | 18517 |
| CL | ml $min^{-1}$ $kg^{-1}$ | 32.5 | 28 | 10.1 | 10.7 | 9.05 |
| $V_{ss}$ | L $kg^{-1}$ | 1.8 | - | 0.56 | - | - |
| $F^b$ | % | - | - | - | - | - |

**Supplementary Table 1 |** Table summarizing pharmacokinetic assessment of dTAG-13 and $dTAG^V$-1 by intraperitoneal (IP) or intravenous (IV) administration.

### SUPPLEMENTARY FIGURES

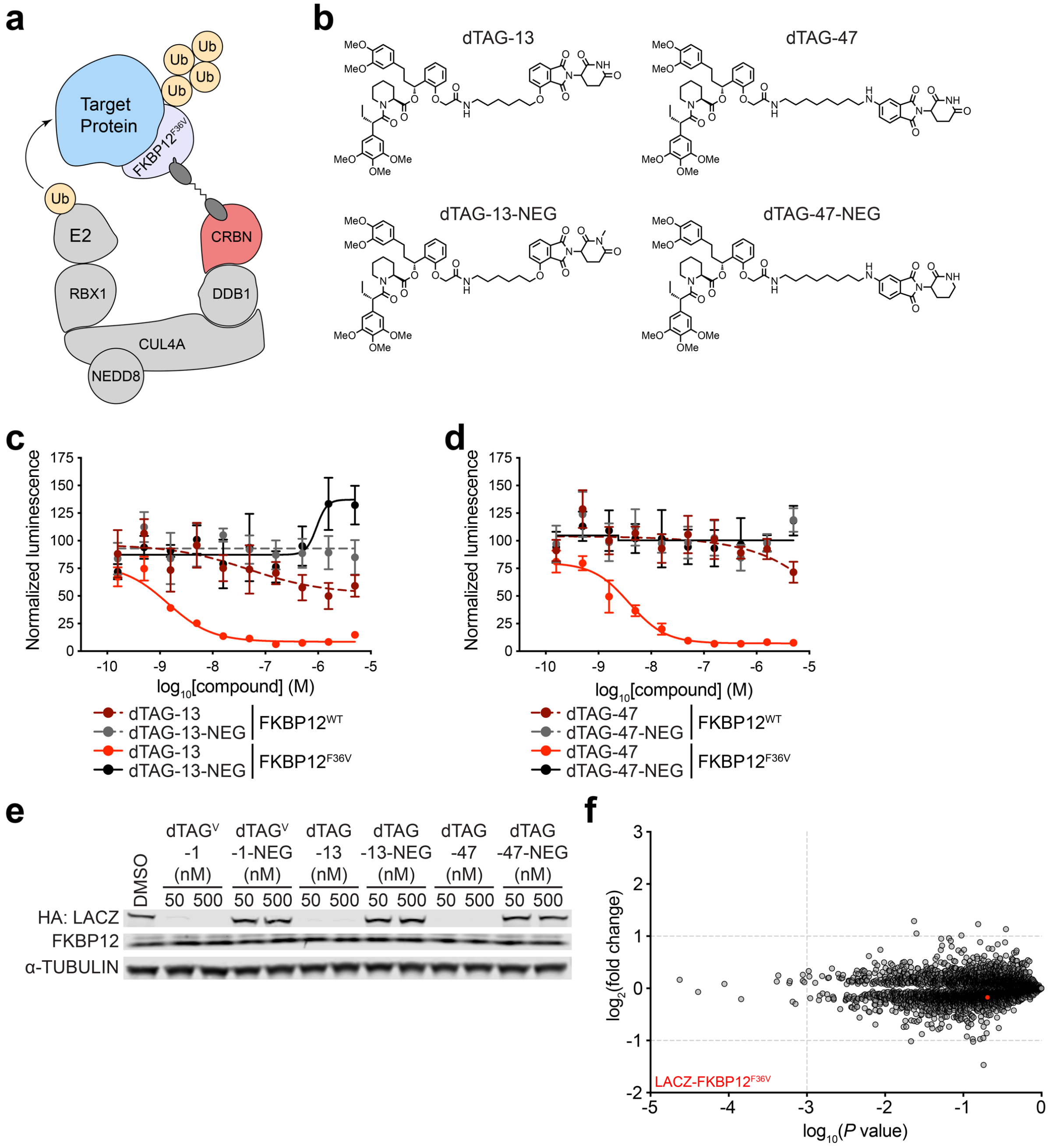


**Supplementary Fig. 1 | Evaluation of CRBN-recruiting dTAG molecules and heterobifunctional control compounds.** (**a**) Schematic depiction of the dTAG system using CRBN-recruiting dTAG molecules. CRBN-recruiting dTAG molecules promote ternary complex formation between the FKBP12$^{F36V}$-tagged target protein and E3 ubiquitin ligase complex, inducing target protein ubiquitination and degradation. (**b**) Chemical structures of dTAG-13, dTAG-13-NEG, dTAG-47, and dTAG-47-NEG. (**c-d**) DMSO-normalized ratio of Nluc/Fluc signal of 293FT FKBP12$^{WT}$-Nluc or FKBP12$^{F36V}$-Nluc cells treated with the indicated dTAG molecules for 24 h. Data in **c-d** presented as mean ± s.d. of $n$ = 4 biologically independent samples and are representative of *n* = 3 independent experiments. (**e**) Immunoblot analysis of PATU-8902 LACZ-FKBP12$^{F36V}$ clone treated with DMSO or the indicated dTAG molecules for 4 h. Data are representative of *n* = 3 independent experiments. (**f**) Protein abundance after treatment of PATU-8902 LACZ-FKBP12$^{F36V}$ clone with 500 nM dTAG$^{V}$-1-NEG for 4 h compared to DMSO treatment. Volcano plots depict fold change abundance relative to DMSO versus *P* value. Significance designations derived from a permutation-based FDR estimation (q < 0.05) are provided in Supplementary Dataset 1. Data are from *n* = 3 biologically independent samples.

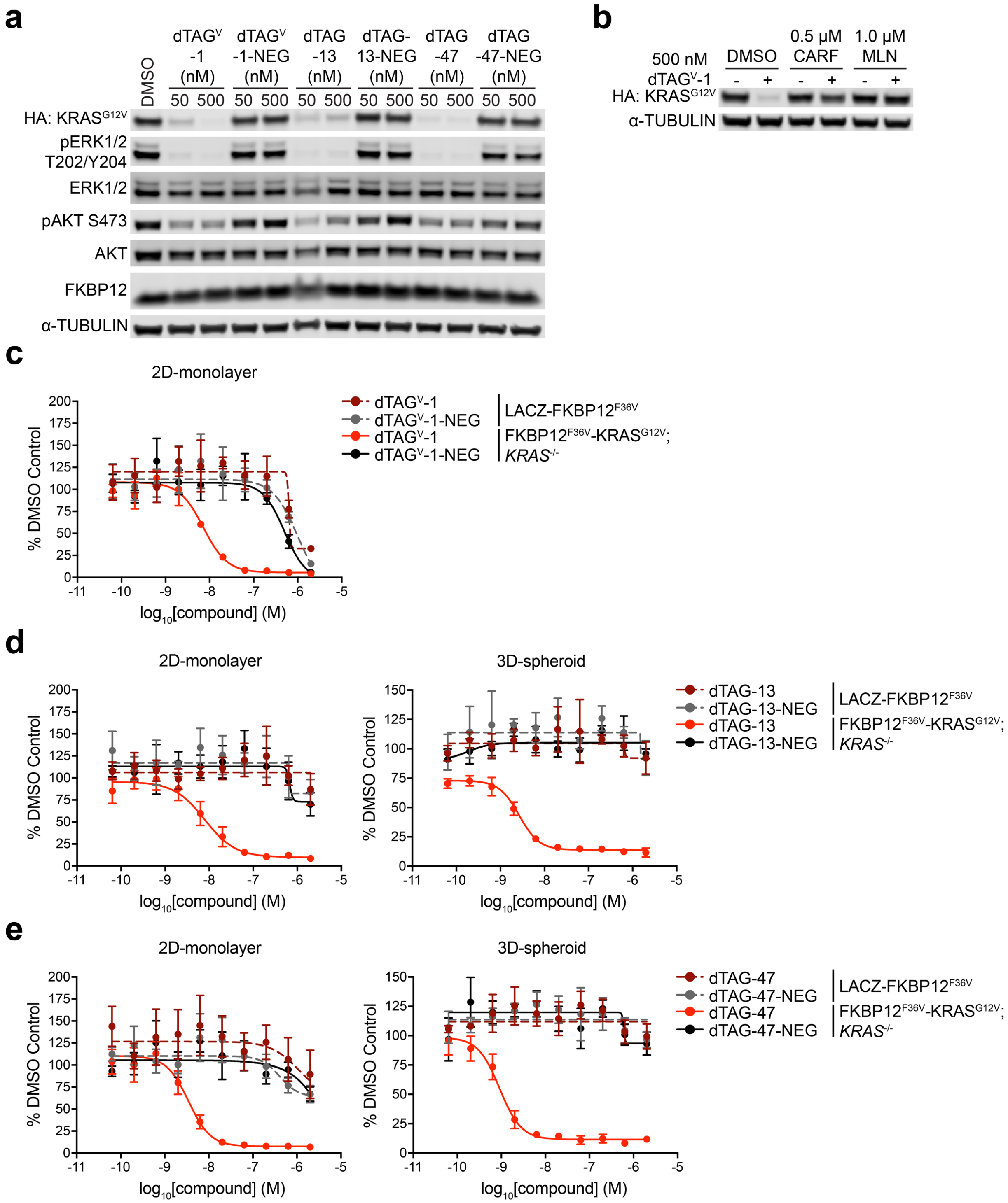


**Supplementary Fig. 2 | Mutant KRAS degradation diminishes aberrant signaling and viability.** (**a**) Immunoblot analysis of PATU-8902 FKBP12$^{F36V}$-KRAS$^{G12V}$; *KRAS*$^{-/-}$ clone treated with DMSO or the indicated dTAG molecules for 4 h. (**b**) Immunoblot analysis of PATU-8902 FKBP12$^{F36V}$-KRAS$^{G12V}$; *KRAS*$^{-/-}$ clone pretreated with DMSO, Carfilzomib (CARF) or MLN4924 (MLN) for 2 h prior to DMSO or dTAG$^{V}$-1 treatment for 4 h. Data in **a-b** are representative of *n* = 3 independent experiments. (**c-e**) DMSO-normalized antiproliferation of PATU-8902 LACZ-

$FKBP12^{F36V}$ or $FKBP12^{F36V}$-$KRAS^{G12V}$; *KRAS*$^{-/-}$ clones treated with the indicated dTAG molecules for 120 h. Cells were cultured as 2D-monolayers or as ultra-low adherent 3D-spheroid suspensions as indicated. Data in **c**-**e** presented as mean ± s.d. of $n$ = 4 biologically independent samples and are representative of $n$ = 3 independent experiments.

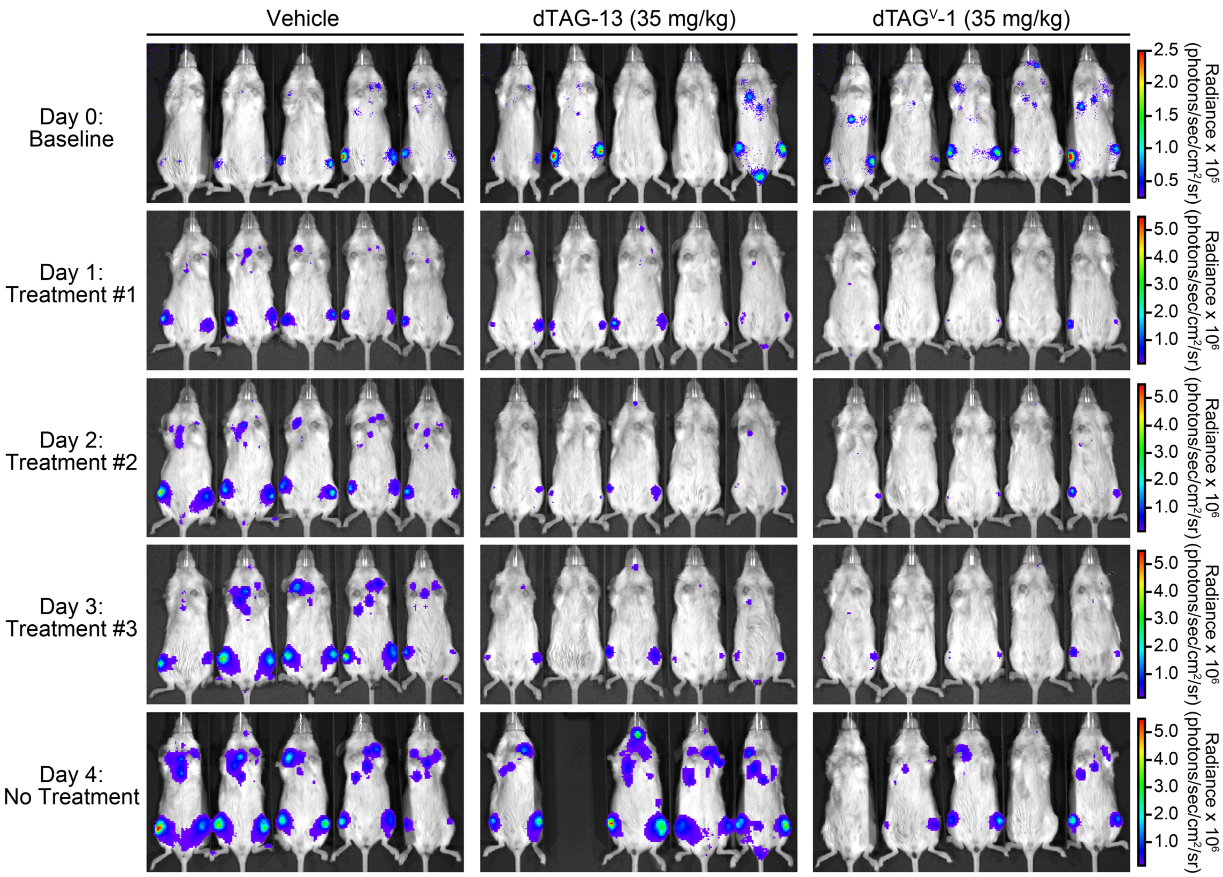


**Supplementary Fig. 3 | dTAG$^V$-1 induces degradation *in vivo*.** Bioluminescence images of vehicle (*n* = 5 biologically independent mice at day 0-4), dTAG-13 (*n* = 5 biologically independent mice at day 0-3; *n* = 4 biologically independent mice at day 4) or dTAG$^V$-1 (*n* = 5 biologically independent mice at day 0-4) treated mice. The same mouse is shown on the same scale at each time point. Quantifications of total flux are provided in Fig. 1i.

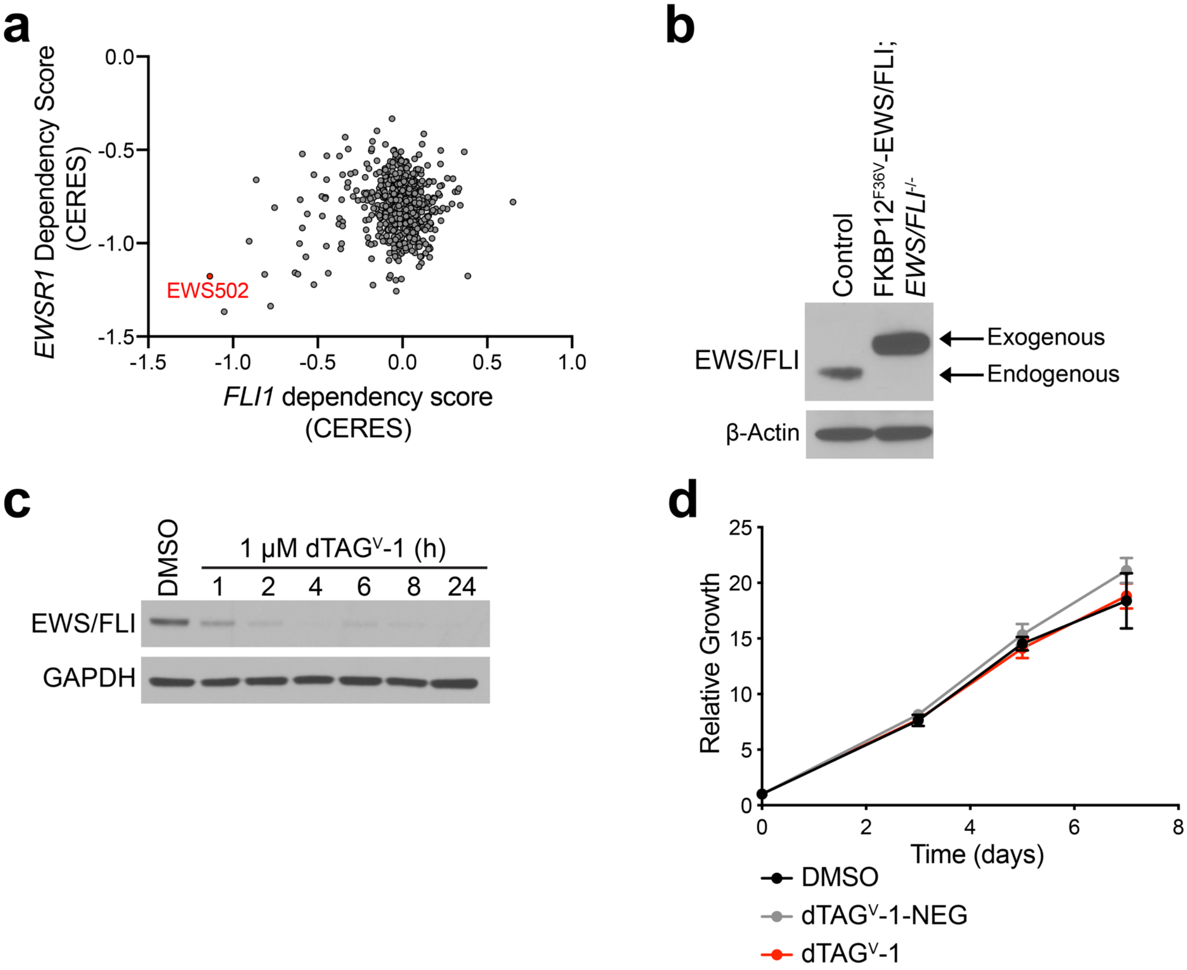


**Supplementary Fig. 4 | Evaluation of degradation with dTAG$^{V}$-1 in Ewing sarcoma cell lines**. (**a**) Dependency score of *EWSR1* and *FLI1* from CRISPR (Avana) datasets from the Cancer Dependency Map portal. (**b**) Immunoblot analysis of EWS502 control and FKBP12$^{F36V}$-EWS/FLI; *EWS/FLI*$^{-/-}$ cells. (**c**) Immunoblot analysis of EWS502 FKBP12$^{F36V}$-EWS/FLI; *EWS/FLI*$^{-/-}$ cells treated with DMSO or dTAG$^{V}$-1 for the indicated time-points. Data in **b-c** are representative of *n* = 2 independent experiments. (**d**) Antiproliferation of EWS502 FKBP12$^{F36V}$-GFP cells treated with DMSO or the indicated dTAG molecules. Y-axis represent luminescence values relative to day 0. Data are presented as mean ± s.d. of *n* = 8 technical replicates and are representative of *n* = 3 independent experiments.

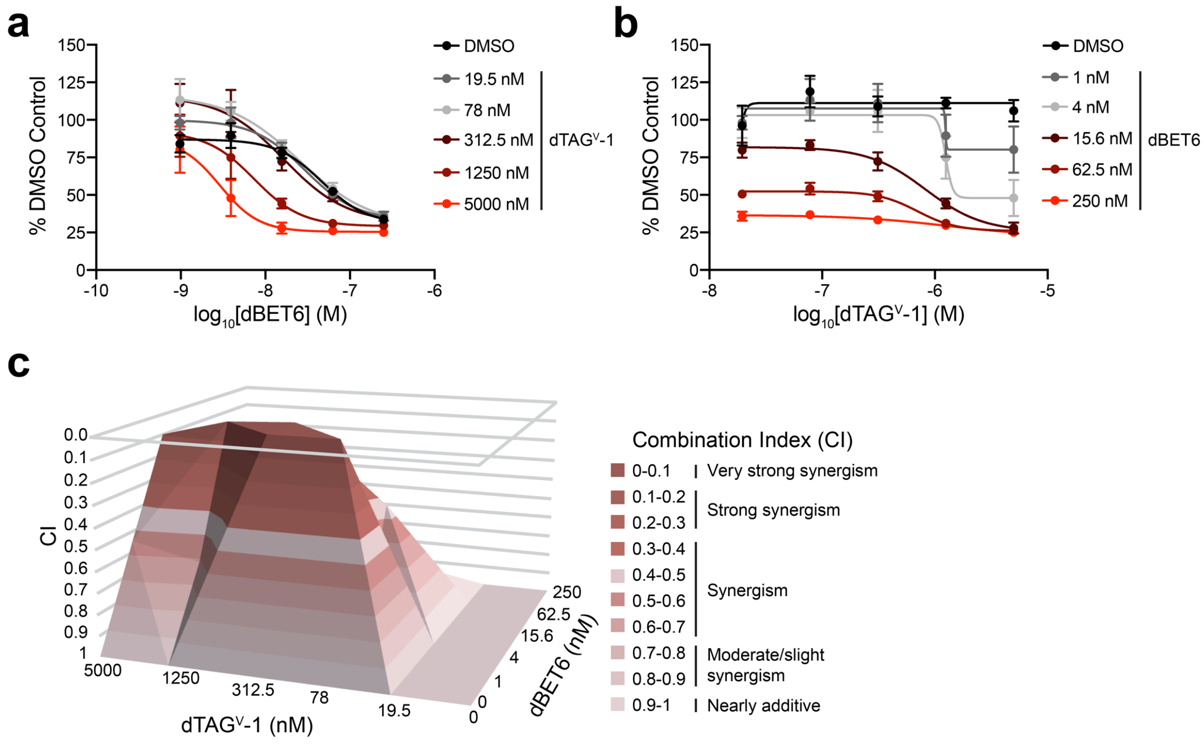


**Supplementary Fig. 5 | EWS/FLI degradation synergizes with BET bromodomain degradation.** (**a-c**) Antiproliferation of EWS502 FKBP12$^{F36V}$-EWS/FLI; *EWS/FLI*$^{-/-}$ cells co-treated with the indicated combinations of dTAG$^{V}$-1 or dBET6 for 72 h. Plots in **a-b** depict DMSO-normalized antiproliferation. Plot in **c** depicts combination index (CI) scores. Data in **a-c** are presented from *n* = 4 biologically independent samples and are representative of *n* = 3 independent experiments.

### SUPPLEMENTARY DATASETS

**Supplementary Dataset 1 |** Mass spectrometry-based proteomics datasets denoting protein ID, fold change, *P* value, and significance are provided for the indicated comparisons in Supplementary dataset 1. A significance designation of '+' refers to $q < 0.05$ derived from a permutation-based FDR calculation.
